# Genomic incompatibilities are persistent barriers when speciation happens with gene flow in Formica ants

**DOI:** 10.1101/2025.03.27.645773

**Authors:** Patrick Heidbreder, Noora Poikela, Pierre Nouhaud, Tuomas Puukko, Konrad Lohse, Jonna Kulmuni

**Affiliations:** Organismal and Evolutionary Biology Research Programme, University of Helsinki, Helsinki, 00560, Finland; Tvärminne Zoological Station, University of Helsinki, Hanko, Finland; Institute for Biodiversity and Ecosystem Dynamics, University of Amsterdam, Amsterdam, 1090 GE, Netherlands; Institute of Evolutionary Biology, University of Edinburgh, Edinburgh, EH9 3JR United Kingdom; CBGP, Univ Montpellier, CIRAD, INRAE, IRD, Institut Agro Montpellier, Montpellier, France

**Keywords:** barrier loci, Bateson-Dobzhansky-Muller Incompatibility (BDMI), network, demographic modeling, speciation

## Abstract

A current goal of speciation research is to identify the loci underlying reproductive barriers between species. Locating such barrier loci in empirical data is difficult due to the often complex demographic history of diverged taxa and the heterogeneity in evolutionary forces across the genome. Here we take advantage of a natural case of hybridization between two wood ant species (*Formica aquilonia* and *F. polyctena*) to identify regions of reduced long-term gene flow using demographically explicit scans of non-admixed genomes. In addition we identify candidate Bateson-Dobzhansky-Muller incompatibilities (BDMIs) through an imbalanced recombinant haplotype frequency analysis of natural *F. aquilonia* × *F. polyctena* hybrid genomes. Both approaches find barriers that are scattered across the genome. Furthermore, candidate BDMIs significantly overlap with the long-term barriers identified by gIMble, indicating that incompatibilities have persisted despite divergence with gene flow between the wood ant species. Intriguingly, BDMIs interact in a network and the number of pairwise interactions a BDMI has correlates with its long-term barrier strength: hub-like BDMIs with many pairwise interactions reduce gene flow more effectively. Finally in regards to function, long-term barriers identified by gIMble arise outside regions of both gene coding sequences (CDS) and transposable elements. In contrast, regions where long-term barriers and BDMIs co-locate are significantly associated with introns, implying a potential role of alternative splicing or gene regulation in incompatibilities, rather than CDS divergence. Overall, our results highlight the underappreciated impact of multilocus BDMIs and the need to consider network connectivity of BDMIs in future work.

**Significance:** Detecting barrier loci that reduce gene flow between closely related species is a common goal of speciation research. However, reliable detection of barrier loci is difficult due to confounding signals in genomic data. Here we take advantage of two different, recently developed approaches and find that barrier loci between wood ant species are scattered across the genome, and despite on-going gene flow, maintain two distinct species. We reveal that genomic regions that are incompatible between the two species can act as persistent barriers, despite theoretical predictions for their collapse under gene flow. Connectivity between incompatibilities also seems to play an important role in barrier persistence. These results highlight the need to consider connectivity between barrier loci in future speciation research.

## Introduction

Current work on the genomics of speciation focues on identifying loci or regions in the genome that underlie reproductive barriers between diverging lineages (1–3). However, how reproductive barriers and their genomic architecture evolve during speciation with gene flow remains an open question. A barrier locus can be defined most broadly as any locus that reduces the flow of neutral alleles between taxa (4, 5) by acting as a barrier to the production of unfit hybrids (1). This could occur either through effects on habitat choice, local adaptation, or assortative mating, or because the barrier locus is involved in negative epistasis that reduces the fitness of hybrids, i.e. Bateson-Dobzhansky-Muller incompatibilities (BDMIs) (6–8).

Detecting barrier loci from genomic data is challenging. A primary method for their detection has been the use of genome scans of summary statistics to identify outlier genomic windows between distinct lineages. Since barrier loci locally reduce gene flow, they should be associated with locally elevated genetic divergence (*d*_xy_) and differentiation (*F*_ST_) in the genome (1, 9). However, a fundamental problem with simple genome scans is that summary statistics are affected by the stochasticity of drift and migration, heterogeneity in selective forces, as well as the genetic (i.e. mutation, recombination) properties of the populations being compared (10). Barrier regions, however, are specifically defined by a reduction in the effective rate of migration or gene flow. Consequently, summary statistics do not allow us to distinguish reduced migration from other population genetic processes that generate elevated divergence or differentiation. Recognition of the limitations of genome scans has driven the development of new methods to infer barrier regions via a signal of reduced effective migration while controlling for confounding factors. These long-term barriers may encompass various pre- and postzygotic mechanisms that contribute to reproductive isolation. Examples of these new methods include DILS, RIDGE (11, 12), and gIMble (13). However, while model-based genome scans can identify regions of reduced gene flow, they are agnostic about the nature of the barrier loci involved and whether barrier loci involve strong epistatic interactions.

After the very initial stages of divergence, we expect multiple reproductive isolating mechanisms to act in parallel, with their associated genomic regions potentially interacting via epistasis or coupling (4, 14–17). To what extent interactions between the reproductive isolating mechanisms and the underlying barrier loci increase barrier strength and promote speciation is a focal question. A fundamental model of epistatic interactions in speciation is the BDMI model. According to this model, alleles that arise in two separate populations have the potential to reduce the fitness of hybrid individuals when combined in a single genome, as these alleles have never coexisted in the same genetic background. BDMIs can be purely intrinsic, meaning they reduce fitness of hybrids independent of the environment (18), or impacted by ecological selection as highlighted in recent work(14, 19). Theoretical work on the basic BDMI model has focused on the number of BDMIs for a given divergence level, the number needed for speciation, and the persistence of BDMIs in populations which exchange migrants (20–25). These studies find that two-locus BDMIs are not persistent barriers under secondary contact and tend to be selected against and removed (26, 27). In contrast, BDMIs that are also under ecological selection (14, 23) or involve multiple epistatically interacting loci (25), can form more persistent barriers to gene flow in the long term. Biologically more realistic incompatibilities, where genes interact in biological pathways and epistasis involves multiple loci are recognized (28–31). However, few empirical studies have investigated how the structure of the network itself impacts how well incompatibilities persist under gene flow.

The detection of BDMIs can be achieved by investigating ancestry patterns in hybrids, either in laboratory crosses or in natural hybrid populations. Early mapping studies have successfully identified numerous BDMIs in various species, including *Drosophila* (32, 33), *Solanum* (34, 35), and house mice (36). However, laboratory crosses are only feasible in suitable model systems and can be laborious to perform. More recently, the X(2) statistical method for detecting BDMIs from whole-genome data in natural hybrid populations has been developed based on detecting an underrepresentation of one of the recombinant genotypes between two loci (37). For instance, using data from three natural hybrid swordtail fish populations, Li et al. (37) identified 54-66 putative incompatibilities, with 0.5% genomic divergence per site (38). However, the extent to which BDMIs contribute to persistent, long-term barriers to gene flow remains an open question, and to date, there are few empirical studies examining genomic barriers across time scales (36, 39). Here we use the term persistent BDMIs to describe BDMIs that remain present between two lineages or species despite gene flow.

We still have little information on what are the general characteristics of barrier loci (i.e. genomic regions with reduced effective migration rates). There are expectations for barrier and BDMI accumulation in regions of reduced recombination (such as within chromosomal inversions or centromeric regions (40, 41)), as observed in certain butterfly species (13, 39, 42, 43) and in gene rich regions (44 but see 39). While the BDMI model itself does not specify the mechanism causing incompatibilities between loci, it is generally expected that BDMIs will arise in functional regions, such as coding regions of genes or transposable element (TEs) insertion sites (45–47). It has also been increasingly recognized that introns and intergenic sequences may play crucial roles in reproductive isolation, particularly through regulatory elements that influence gene expression (47, 48). For example, aberrant splicing may contribute to reproductive isolation by reducing hybrid fitness, with spliceosome genes acting as BDMIs (49). One aspect of barriers and BDMIs that has received less attention in empirical work is the potential non-independence of barrier loci. Theoretical work highlights how coupling or epistasis can facilitate the build-up of strong reproductive isolation (25, 50). Here we extend this to BDMIs, investigating if multi-locus BDMIs can act as persistent barriers even with gene flow.

Red wood ants of the *Formica rufa* (Hymenoptera: Formicidae) group serve as a good model to investigate the nature of barriers to gene flow for several reasons. First, this group contains several recently diverged lineages that still hybridize in their natural habitat. Despite high rates of hybridization, the species remain distinct in sympatry (51). Second, ants are arrhenotokous haplodiploids (52), meaning that fertilized eggs develop into diploid females and unfertilized eggs develop into haploid males. Given their haploidy, hybrid males can be used to map both recessive and dominant incompatibilities genome-wide (53). Here, we focus on two species, *F. aquilonia* and *F. polyctena*. Both are polygynous, with hundreds of reproductive queens within a nest, resulting in low relatedness among nestmates, and polydomous, with populations consisting of networks of interconnected nests. Polygyny in *Formica* is controlled by a set of three inversions on chromosome 3, called the social chromosome (54), and even though both species are polygynous, they have different forms of the supergene haplotype, generating the potential of barrier enrichment in the social chromosome (55). A previous reconstruction of the speciation history suggests that the two species diverged around 500 kya (56, 57), with unidirectional gene flow from *F. aquilonia* into *F. polyctena* (57). Several stable natural hybrid populations exist, one of which has been studied for 20 years in southern Finland (58, 59). Based on earlier findings, hybrid incompatibilities segregate in this hybrid population (58, 60).

Here, we use the *Formica* ant system to identify long-term barriers to gene flow from parental genomes and segregating BDMIs from a young hybrid population (Fig. 1), and ask the following questions. First, how are long-term barriers to gene flow distributed across the genome? In the face of gene flow, barrier loci may be expected to become clustered into a few genomic regions, which are easier to maintain by selection than scattered barrier architectures. Second, is there an overlap between the long-term barriers and BDMIs, suggesting that persistent BDMIs can evolve as strong barriers even in the presence of gene flow? Given the different time scales of the barriers and the fact that long-term barriers may encompass both pre- and postzygotic barriers, whereas BDMIs are solely associated with postzygotic isolation and may easily collapse under gene flow, we expect some but not complete overlap between the different types of barriers. Third, are BDMIs that involve a large number of loci associated with a stronger reduction in long-term gene flow than pairwise BDMIs? We hypothesize that multi-locus incompatibilities with more epistatic interactions across the genome are harder to resolve and thus expected to resist gene flow better than simple incompatibilities. Finally, do long-term barriers and BDMIs concentrate in regions of low recombination, and are they associated with CDS, introns or TEs?

**Figure 1.**
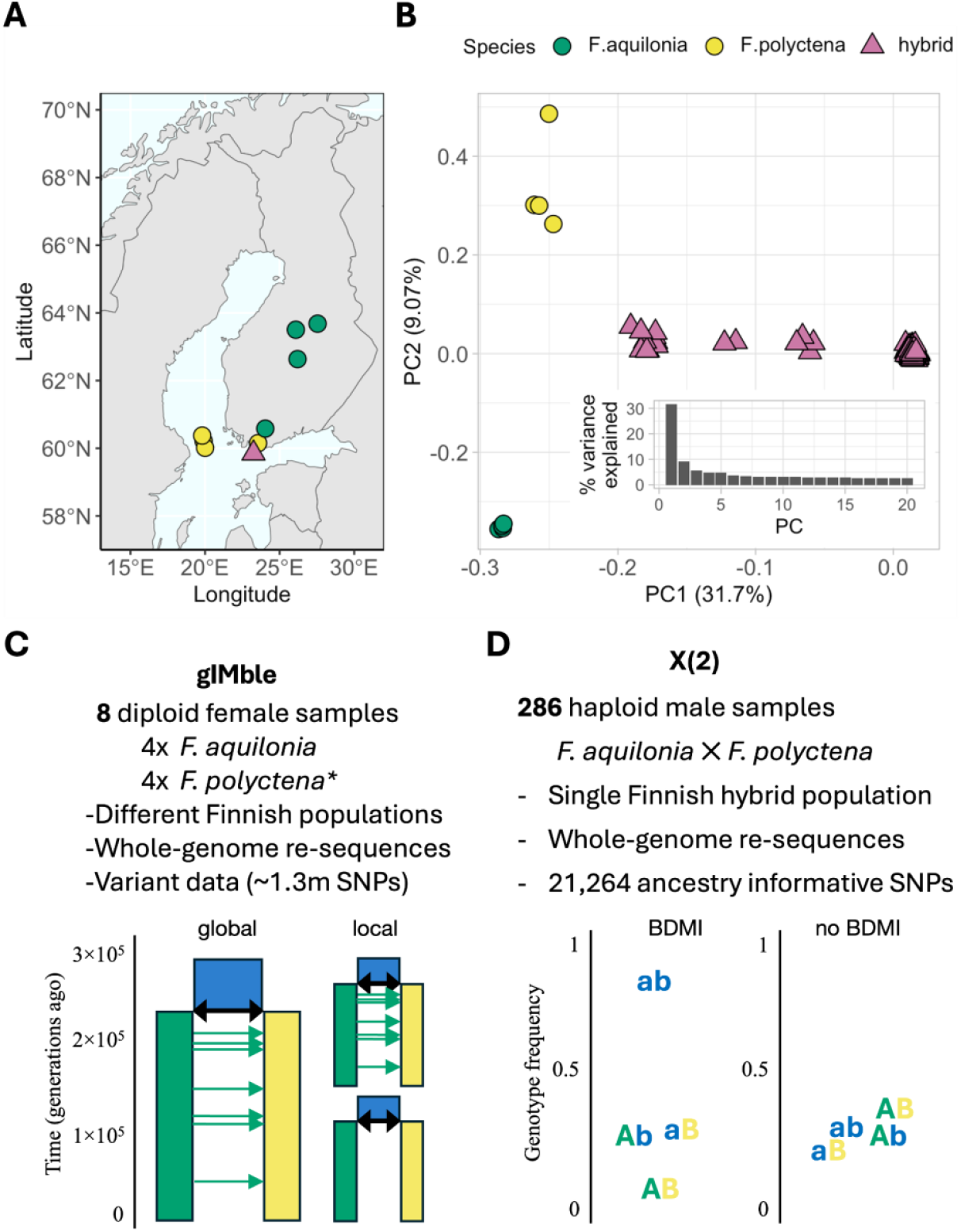
Sampling data and methods used for detecting barrier and incompatibility loci. (A) Sampling locations of *F. polyctena, F. aquilonia*, and their hybrids. Green and yellow circles respectively indicate *F. aquilonia* and *F. polyctena* individuals sampled from their sympatric range (8 diploid individuals altogether), purple triangles indicate the Långholmen population where 286 haploid hybrid individuals were sampled. (B) PCA of samples used to detect barrier regions from the genome (symbols as in panel A). Inset barchart is the variance explained by principal components 1-20. (C) Data used for identifying long term barriers with gIMble (13) by comparing a local demographic model of divergence with gene flow (green arrows) in a genomic window against a global genome-wide demographic model. Barriers are detected when a window has significantly less gene flow than the global model. (D) Data used for identifying BDMIs within a natural hybrid population using X(2). A BDMI is inferred when one of the recombinant genotypes (AB) is underrepresented in contrast to parental genotypes (Ab, aB) and the alternative recombinant (ab). Uppercase A and B represent derived alleles in the green and yellow lineages, while blue lowercase a and b represent the ancestral alleles.

## Results

### Divergence with gene flow has resulted in barriers that are scattered genome-wide

We used two complementary methods and datasets to map barriers to gene flow. First, we identified long-term barriers between *F. aquilonia* and *F. polyctena* throughout their divergence history using whole-genome resequencing (WGS) data from four samples of each species with gIMble (v1.0.3) (13)(Fig. 1A-B). This method accounts for demographic history and models barriers as heterogeneity in effective migration rate (*m*_*e*_) (Fig. 1C) in sliding windows (median length 41.5kb; Fig. S1) across the genome. The underlying model assumes unidirectional post-divergence gene flow from *F. aquilonia* into *F. polyctena* (1.5 migrants per generation, ∼600 kya; Fig. S2, Table S1, Supporting text), consistent with previous findings (57). The estimates of *N*_*e*_ (*N*_*F*. *aqu*_, *N*_*F*. *pol*_, *N*_anc_) and *m*_e_ vary considerably along the genome (Fig. S3). Local support for barriers to gene flow were defined as Δ_B0_>0, where a history of reduced *m*_*e*_ fits better than a model assuming the global *m*_*e*_ estimate of 5.19E-06, which translates into 1.50 individuals per generation (Table S1).

Overall, we found that 5.3% (1,110 out of 21,042) of genomic windows showed Δ_B0_ > 0, indicating potential barriers to gene flow (Fig. S4). However, after accounting for a FPR of 5%, 41.5% of these windows (461 out of 1,110) were excluded as potential false positives (Fig. S4). By merging overlapping windows in the remaining set of 649 windows, we identified 283 distinct candidate barrier regions, spanning a total length of 17.4Mb, which constitutes 8.2% of the genome (Fig. 2A-B, Fig. S4). These barrier regions varied in length from 33.3kb to 215.1kb, with a median length of 51.4kb. The barriers were not enriched on specific chromosomes but were instead distributed across all chromosomes, each containing approximately the same fraction of barriers (Fig. 2A, Fig. S4, Table S2). Notably, the strongest barriers to gene flow only partially overlapped with F_st_ and d_xy_ outliers (Fig. S5), demonstrating the importance of identifying candidate barrier regions by explicitly modeling variation in *m*_*e*_. Mean and window-wise heterozygosity (*H*), d_xy_ and F_st_ are shown in Table S3 and Fig. S6.

**Figure 2.**
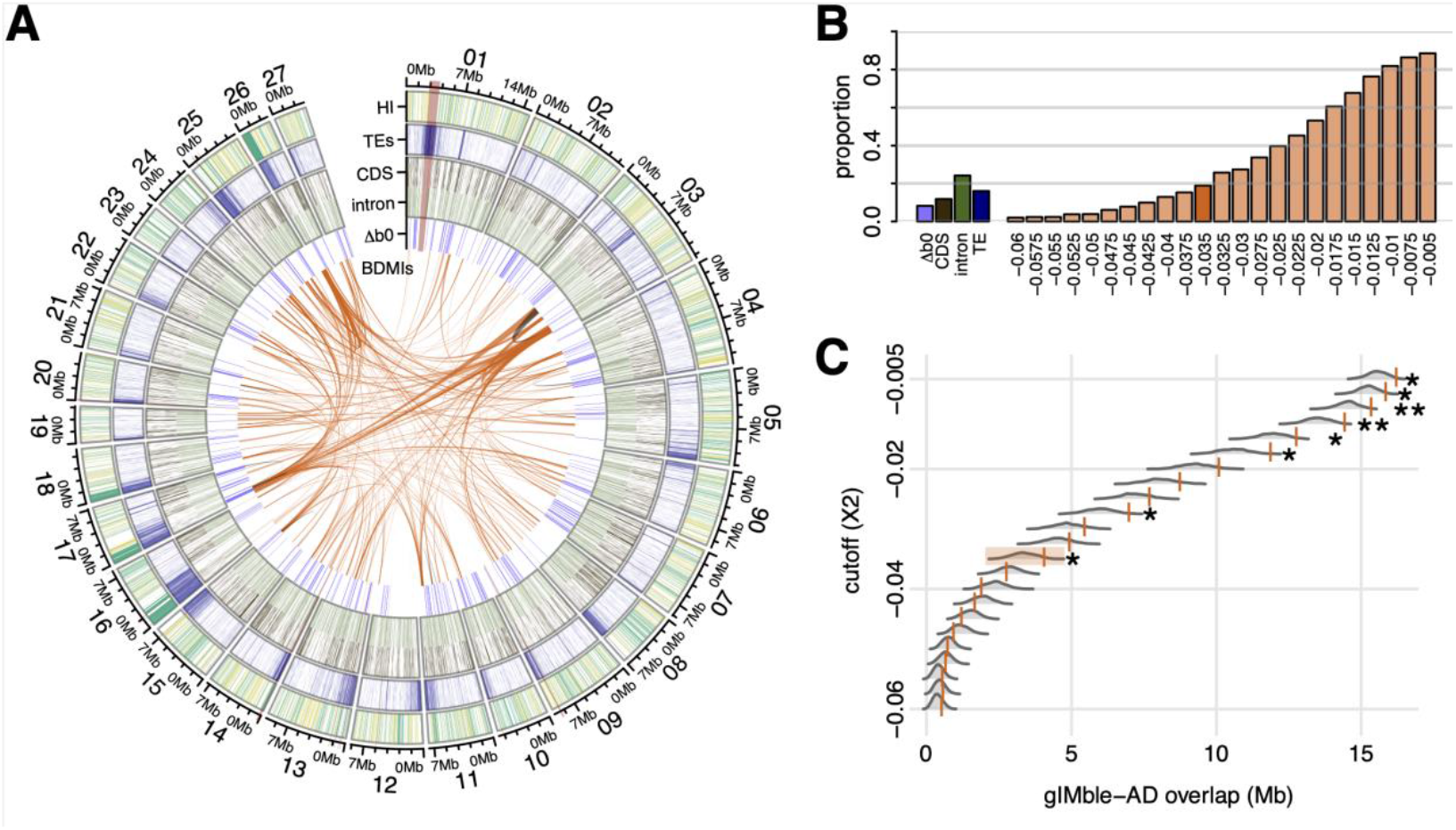
Candidate barriers are genome-wide despite speciation with gene flow. (A) Circular plot showing the locations of genetic incompatibilities (BDMIs), long-term barriers to gene flow (gIMble; Δ_B0_>0 and FPR≤0.05), introns and coding sequences (CDS), TEs, and hybrid index (HI) of sorted genomic regions. Interchromosomal BDMIs are indicated in orange and intrachromosomal in black for X(2) = -0.035 (highlighted in B and C). The inner ring shows long-term barriers in blue. Moving outwards, the rings indicate introns and CDS, TEs and HI. The outermost values represent the 27 scaffolds with centromeric regions highlighted across tracks in red. (B) Percent coverage of the reference genome for each dataset and X(2) threshold. (C) Overlap between BDMIs and long term barriers. Observed values for each X(2) threshold are plotted as orange lines, bootstrapped distribution values are visualized behind the observed values as grey density plots. Overlaps are marked as significant as following: *p<0.05, **p<0.01, ***p<0.001.

In addition to using a demographically explicit barrier scan, we analysed 286 hybrid haploid male genomes from a contemporary *F. aquilonia* × *F. polyctena* hybrid population in Långholmen, Finland, to identify BDMIs (Fig. 1A, Table S4). We did this through an imbalanced recombinant haplotype frequency analysis which identifies two-locus heterospecific allele combinations that occur less often than expected in the absence of selection, indicative of negative epistasis, i.e. BDMIs between parental alleles, through the X(2) statistic (Fig. 1 D)(37).

To identify incompatibilities, a total of 21,283 biallelic Ancestry Informative Markers (AIMs) were used (Fig. S7), which had an allele frequency difference ≥0.95 between parental populations of *F. aquilonia* and *F. polyctena* (12 diploid individuals per species) (59). Specifically, we identified pairs of AIMs in negative LD as quantified by *X*(2) < -0.005 and *D*′ < 0 (37). While the -0.005 X(2) threshold was determined through simulations by Li et al. (37) to discriminate neutral from BDMI loci, X(2) becomes more negative closer to BDMI loci. However, since any X(2) threshold is arbitrary, we explored the properties of a range of increasingly strict thresholds. A sliding window analysis (20 consecutive AIMs per window, windows slide by 4 AIMs, min window size 3kb, max window size 1.67Mb, mean window size 184kb), identified 156,119 window pairs as candidate BDMIs at the X(2) < -0.005 threshold (1.17% out of the 13,325,703 possible window comparisons). Overlapping windows were merged, and merged windows were divided based on mean X(2) into 23 datasets of decreasing X(2) strictness (Fig. S8). The number of candidate BDMI pairs identified across the 23 X(2) thresholds ranged from 87 to 11,486 (Fig. 2A, Table S5). We have used the X(2) < -0.035 dataset throughout the manuscript for visualization, other thresholds are available in the supplementary materials. The sizes of BDMI loci ranged from 3.0kb to 1.7Mb (mean 154kb, median 116kb, Table S5), and covered a total span from 4.53Mb to 188.5Mb (1.9 - 88.5%) (Fig. 2B, Fig. S9). Like the long-term barriers identified by gIMble, candidate BDMIs are scattered throughout the genome. To understand recombination and barriers to gene flow, we used two proxies for areas of low recombination. First, we tested if BDMIs were disproportionately located on the social chromosome characterized by large inversions (54), but found no significant enrichment of BDMIs on the social chromosome (Table S6). Second, we calculated the mean distance from BDMI and gIMble barriers to centromeric regions, finding that both BDMIs and barriers are closer to centromeres than expected by chance (Table S7). Additionally, while chromosome 17 contains no known inversions, we found it had consistently more BDMIs than the genome wide average across all X(2) thresholds (Table S6). Finally, we primarily detected interchromosomal BDMIs across X(2) thresholds (mean interchromosomal proportion 0.96%; Table S5).

### Incompatibilities act as persistent long-term barriers and are characterized by multi-locus interactions

To evaluate the role of BDMIs as persistent long-term barriers to gene flow, we investigated i) whether the two types of barriers overlapped, and ii) whether the number of interactions a BDMI locus has affected the effective migration rate (*m*_*e*_). We found a significant overlap between long-term barriers—identified as regions of reduced *m*_*e*_ between parental genomes using gIMble—and BDMIs detected in hybrids for 8 out of 23 of the X(2) threshold values (Fig. 2C). While total overlap between the two types of barriers ranged from 0.54-16.2Mb depending on the X(2) threshold value (Table S8), the overlaps with the strictest X(2) thresholds (< -0.0425) were not significant.

We analyzed candidate BDMIs as a network (Fig. 3A-B, Fig. S10), with loci as nodes and inferred BDMIs as edges, thereby allowing us to analyze the effect of network topology on barrier strength (Fig. 3A, C). We found that, across all but two X(2) thresholds, the more connected (higher degree) the BDMI locus the stronger the reduction in gene flow (Fig. 3D, Fig. S11). Furthermore, we find that the degree distribution of the observed BDMI networks is much broader than the expected degree distribution of the classic BDMI model as extended by Orr (20)(Fig. 3B, E). In other words, there are hub-like BDMIs in the observed incompatibility network that interact with large numbers of other BDMI loci. These hubs are not found in a network drawn from the Orr model. Bootstrapping 1000 networks revealed further differences across multiple aspects of network topology including max observed node degree, mean path length between nodes, and the average number of neighbors a node has (Fig. 3F-H, Fig. S12). Together, these results suggest that multi-locus BDMIs do evolve, and that network topology may play a role in their maintenance in the face of continuous gene flow between the *Formica* species.

**Figure 3.**
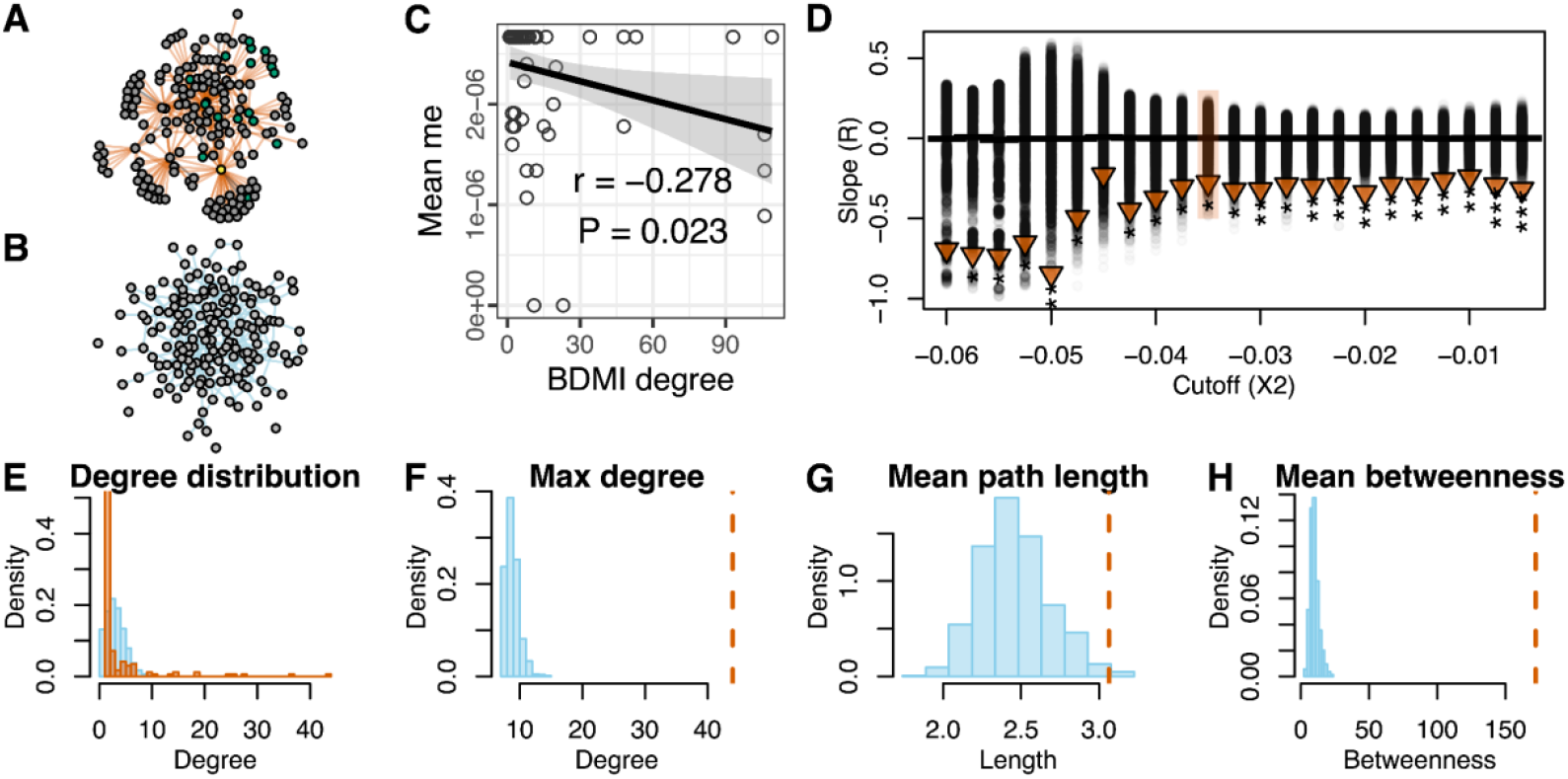
There is connectivity between candidate BDMI loci which impacts how effectively the locus reduces gene flow. (A) Candidate BDMI regions form a network with many hub-like interactions. Network for candidate BDMIs at X(2) = -0.035. (B) Visualization of a BDMI network generated by Orr’s (20) model with the same average node degree as the observed BDMI network. (C) Long-term barrier strength (Δ_B0_) negatively correlates with the degree (number of inferred BDMIs) of BDMI regions at X(2) = -0.035. (D) There is negative correlation between Δ_B0_ and degree for all X(2) values, and only two of the X(2) thresholds (−0.06, -0.045) are not significant. Observed correlation coefficients are plotted as triangles above the bootstrapped distribution of correlation coefficients in black circles. Black horizontal bars represent mean bootstrap correlation coefficients. Observed correlation coefficients are marked as significant as follows: *p<0.05, **p<0.01, ***p<0.001. X(2) = -0.035 is highlighted in orange. (E) Degree distribution of the observed BDMI network in orange, against the degree distribution generated by the Orr model in blue. (F-H) Observed BDMI network values (orange dashed lines) versus values from 1000 bootstrapped Orr networks (blue histograms). Max degree is the highest number of edges that are connected to the same node. Mean path length is the average number of edges to travel between two random nodes. Mean betweenness is the average number of paths through each node.

### Long-term barriers, BDMIs and their overlap show contrasted association with functional elements and ancestry sorting

We investigated the association between barrier loci and functional elements, including coding sequences (CDS), introns, and TEs, to investigate their role in barrier loci. Long-term barriers (Δ_B0_>0) overlapped significantly less with CDS and TEs than expected by chance (circular resampling p<0.001, Fig. 4A). In contrast, BDMIs were significantly associated with CDS and introns (Fig. 4A) for four and two X(2) thresholds, respectively. The genome-wide association between BDMIs and TEs was strongly dependent on X(2) thresholds: BDMI regions defined with strict X(2) (≤ -0.035) thresholds were significantly enriched for TEs, while those defined with the least extreme X(2) thresholds (≥ -0.0125) showed a depletion for TEs (Fig. 4A). Finally, persistent BDMIs— regions where candidate BDMIs overlapped with long-term barriers—were depleted for CDS and TEs for the least strict X(2) thresholds, but showed significant enrichment for introns for the most strict X(2) thresholds (Fig. 4B). Long-term barriers, BDMIs, and their intersections are associated with different functional features, suggesting a complex network involving coding regions, regulatory motifs, and a potential role for transposon activity in the barriers between species.

**Figure 4.**
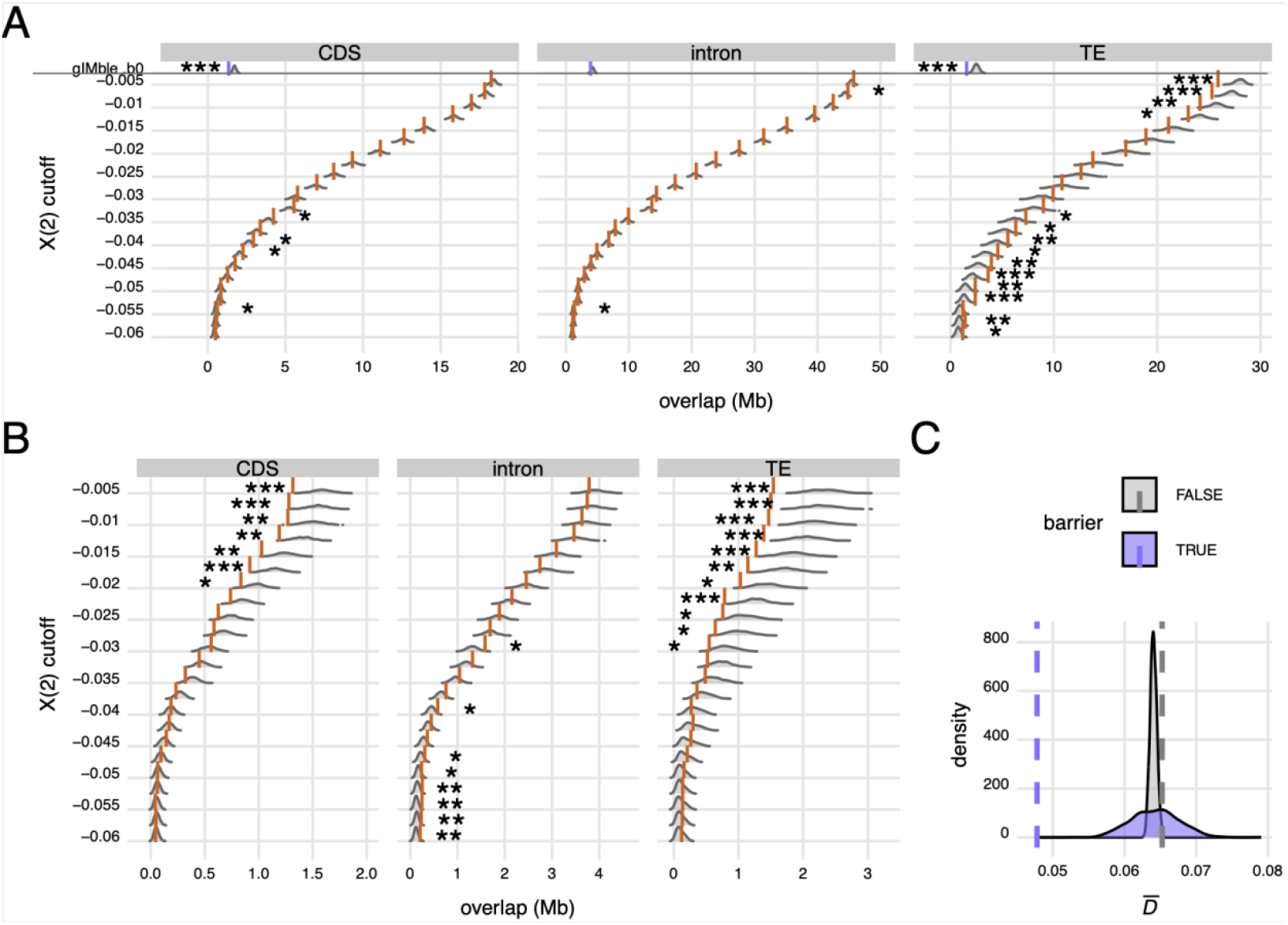
Functional features associated with barriers. (A) Overlap between long-term barriers (b0, highlighted) or BDMIs (with various strictness of X(2) thresholds) and coding sequences (CDS), introns and TEs. (B) Overlap between long-term BDMIs (regions shared by long-term barriers and BDMIs) and CDS, introns and TEs. Observed values for b0 and each X(2) threshold are plotted as orange lines, with bootstrapped distribution values displayed as grey density plots in the background. Overlaps are marked as significant as follows: *p<0.05, **p<0.01, ***p<0.001. (C) Observed 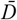 in barrier windows (purple vertical dashed line) is significantly less than in non-barrier windows (grey vertical dashed line and grey density distribution; circular resampling, p < 0.001) and lower than expected by chance if barriers were randomly distributed (purple density distribution; circular resampling, p < 0.001).

Finally, we identified a total of 13 genes in persistent BDMIs where there is no gene flow between species (Table S9). Most of these are involved in core cell functions that have been found disrupted by BDMIs identified in other taxa. Five of the genes within persistent BDMI regions are implicated in the development of cancer, aligning with the findings in swordtail fish, where one of the mapped incompatibilities causes melanoma in hybrids (61). Furthermore, two genes were ribosomal and one involved in maintaining mitochondrial function, all candidate functions that have been associated with incompatibilities in other studies (62–65) (Table S9). These genes may have contributed to the establishment of strong BDMIs, which have effectively reduced gene flow between *F. aquilonia* and *F. polyctena* during their evolutionary history and formed a barrier that has not collapsed despite gene flow.

To understand how ancestry has sorted over time in hybrid genomes we calculated 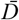 (66) as the variance in HI in gIMble barrier and non-barrier regions. 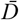 indicates sorted ancestry within regions when close to zero and individual HI are around 0.5. We found that in the hybrid population, 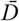 in gIMble barrier regions (Δ_B0_>0) is significantly lower than in non barrier regions (circular resampling, p < 0.001)(Fig. 4C).

## Discussion

One of the central goals in evolutionary biology is to identify the genetic basis of reproductive isolation. Little is known about whether long-term barriers to gene flow and intrinsic barriers observed in hybrids involve the same loci and genomic regions. Theoretical models of speciation have shown that pairwise incompatibilities are unlikely to persist in the presence of gene flow (23). In contrast to these models, we find a significant overlap between BDMI regions and long-term barriers between *F. aquilonia* and *F. polyctena*, suggesting that BDMIs have effectively reduced long-term gene flow between these ant species. Intriguingly, we find that the number of pairwise interactions a BDMI has correlates with its long-term barrier strength: BDMIs with many pairwise interactions to other BDMIs reduce gene flow more effectively than those which have fewer interactions. These results highlight the underappreciated impact of multilocus BDMIs as barriers for gene flow.

The two wood ant species *F. aquilonia* and *F. polyctena* diverged approximately half a million years ago, exchanging on average 1.5 migrants per generation after their divergence. Both divergence time and migration rate estimated here are consistent with previous findings in these species (57). Between sympatric populations of *F. aquilonia* and *F. polyctena*, we find that about 8% of the genome acts as persistent long-term barriers to gene flow. Instead of being localized to a few regions of the genome, as for example seen in *Heliconius* butterflies, these long-term barriers are scattered throughout the *Formica* genome, supporting the growing body of evidence for polygenic barriers to gene flow (26, 39, 67–69). We find that for a wide range of X(2) cut-offs, BDMIs are also scattered throughout the genome. While co-expressed genes often cluster on the same chromosome (70), and most physical protein interactions are intrachromosomal (71), we did not find any evidence for an enrichment of intrachromosomal BDMIs, in agreement with a previous BDMI scan in swordtail fish (37). Unlike our definition of long-term barriers, the screen for BDMIs relies on arbitrary thresholds. This makes a direct comparison of the number of BDMIs identified to other systems or theoretical expectations difficult.

To date, only a few studies have quantified genetic barriers across multiple time scales. A recent meta analysis by Frayer & Payseur (72) compared incompatibilities and barrier loci identified in lab crosses with barrier estimates from a natural hybrid zone in the house mouse, finding no significant overlap. In contrast, Ebdon et al. (39) observed a significant overlap between long-term barriers, characterized by reduced effective migration rate, and barriers in a natural hybrid zone in *Iphiclides* butterflies. In the current study, BDMIs overlapped significantly with long-term barrier regions across multiple X(2) thresholds, suggesting that some BDMIs contributed to long-term barriers between the two *Formica* species and have persisted in the face of gene flow. We would expect any overlap between long-term and current barrier regions to be incomplete for two reasons. First, long-term barriers may be associated with both postzygotic and prezygotic isolating mechanisms, unlike BDMIs, which are usually thought to involve intrinsic postzygotic mechanisms such as incompatible protein interactions (18, 73). For example, long-term barriers in *F. aquilonia* and *F. polyctena* might contribute to habitat isolation and temporal isolation since the species have different ecological niches and the parental species have different mating flight times (74–76). Second, owing to divergence with gene flow, many emerging BDMIs are constantly being selected against and removed (26, 27), particularly if they involve recessive alleles which are exposed to purging selection in haploid males. Furthermore, the removal of strongly deleterious BDMIs during the establishment of our focal hybrid population may explain why we find no significant overlap between BDMIs and long-term barriers for very stringent X(2) thresholds.

The BDMIs that were found to most effectively reduce long-term gene flow had two distinct and surprising features: i) they were multilocus, i.e. involving more pairwise interactions than expected by chance for a range of X(2) thresholds and ii) they were significantly associated with introns but not with CDS. While the simplest BDMI model involves two loci and two alleles (20, 22), such pairwise incompatibilities are not expected to be able to contribute to long-term barriers in the face of gene flow, as they can be easily purged following introgression (23). In contrast, the complexity of a BDMI increases with the number of alleles (3) or loci involved (25, 77), and BDMIs involving epistasis between three or more loci can act as strong barriers even with gene flow (25). However, these theoretical expectations have not been tested in empirical data. Here we use the number of pairwise interactions that each candidate BDMI locus is involved in (i.e. the BDMI degree) as a proxy for the complexity of incompatibilities. While we do not test for three-way interactions explicitly, trigenic interactions have been shown to often overlap with digenic interactions in protein-protein interaction networks in yeast (78). Thus, our finding that the more pairwise interactions a BDMI has, the more effectively it reduces gene flow, suggests that BDMI complexity determines the long-term maintenance of a BDMI. We note that in the limit, a locus that is involved in a large number of negative epistatic interactions, becomes incompatible with the entire genetic background and it becomes nonsensical to attempt to identify individual incompatibilities. Furthermore, the network of BDMIs we observe is much more connected than predicted by classic null models of incompatibility accumulation (20). Our results, together with other recent studies (29, 79, 80) highlight the need to consider more realistic network models in future BDMI work. While the association between BDMIs, CDS and TE regions may not be surprising, the fact that persistent BDMIs have overrepresentation of intronic sequences is. These results may indicate that BDMIs that most strongly resist gene flow involve regulatory elements that affect gene expression (47, 48). Alternatively, they may simply reflect a survivorship bias; if BDMIs in coding sequences are more deleterious, they would be purged more quickly compared to intronic BDMIs (81, 82).

Barrier regions also appear to influence hybrid genome evolution. The finding that the mean variance in HI 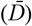 in barrier regions is smaller than in non barrier regions implies both that sorting to single ancestry has occurred faster in barrier regions than outside of them, and that sorting within regions occurs in the same direction. One other study examining 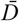 in a *Heliconius* hybrid zone found that 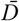 is higher in hybrids (39). However this is not surprising as homozygous samples from either side of a hybrid zone are maximally unsorted 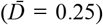. The Längholmen hybrid population, however, is ∼40 generations old, which has allowed time for ancestries to mix and sort in hybrid genomes. Consistently sorting to single ancestry within regions across samples agree with previous results in the *Formica* system which used 19 samples (59).

While studies that map incompatibilities directly using laboratory crossing experiments in model organisms have limited resolution (72, 83), indirect scans for BDMIs based on population genomic data from natural hybrid populations suggest that testing for non-random associations between epistatically interacting alleles (37) poses other challenges: The fact that all pairwise interactions must be examined (in this study we had 13,325,703 possible window comparisons), means that multiple testing leads to difficulty in identifying significant outliers (77). Embracing this complexity, we have conducted all analyses for 23 subsets of candidate BDMIs, defined using increasingly strict (more negative) X(2) thresholds which correspond to sets of BDMI pairs between regions with increasingly distorted ancestry combinations (Fig. S13). The main limitation of this approach is that we cannot meaningfully estimate the absolute number of BDMIs between the two species. Thus an obvious avenue for future research will be to obtain quantitative predictions for both the number and complexity of BDMIs and the LD structure associated with them in samples from contemporary hybrid populations. Another promising future direction is to intersect BDMI and barrier scans with direct estimates of the recombination map and structural variants (which is not yet available for *Formica* ants) to test to what extent barriers and BDMIs accumulate in regions of low recombination and recombination modifiers such as inversions. Our two proxies for low recombination, the social chromosome with know inversions, and distance from barriers to centromere region gave conflicting results with no enrichment in inversions (Table S6), but both candidate BDMIs and long-term barriers being closer to centromeres than expected by chance (Table S7), suggesting future investigations are promising.

In conclusion, our overlay of long-term barriers to gene flow and BDMIs in a hybrid population in the wood ant system demonstrates that – contrary to predictions from simple null models of intrinsic incompatibilities – BDMIs (in particular those involving multiple loci) can act as persistent barriers to gene flow. The enrichment for intronic sequence we find in persistent BDMI suggests a potential role of regulatory elements in the evolution of species barriers. Our study opens the door to future work that compares gene expression and alternative splicing in persistent BDMIs between hybrids and parental species and relates the observed distribution of incompatibilities (both in terms of genomic location and complexity) to explicit predictions based on gene networks.

## Materials and Methods

A summary of the materials and methods is provided here, with detailed information available in the Supporting Information. Briefly, we used two complementary methods and datasets to identify barriers to gene flow. First, we analyzed whole-genome sequencing (WGS) data from four samples each of sympatric *F. aquilonia* and *F. polyctena* from Finland (Fig. 1) using gIMble (v1.0.3; (13)). This method identified long-term barriers by accounting for demographic history and variation in effective population sizes, revealing genomic regions of reduced migration. The model assumed unidirectional gene flow from *F. aquilonia* to *F. polyctena* (Table S1, Fig. S2), consistent with previous findings (57). Local support for barriers was defined as Δ_B0_ > 0, and false positive rates (FPR) were estimated using a parametric bootstrap. Second, we analyzed 286 haploid male hybrid genomes from a single *F. aquilonia* × *F. polyctena* hybrid population to identify BDMIs. This was achieved through an imbalanced recombinant haplotype frequency analysis, which detects negative epistasis between loci in hybrid populations quantified by the X(2) statistic from Li et al. (37). The Långholmen hybrid population in southern Finland, being 24-50 generations old with equal admixture proportions, provided an ideal study system (51). We investigated the connectivity of the candidate BDMIs by calculating the number of BDMI interactions as each locus, and calculated network measures with igraph in R. Finally, to investigate the association between long-term barriers and BDMIs and to quantify the overlap of each with functional genomic partitions (CDS, introns, TEs) we used a circularised data resampling procedure which involves circularising chromosomes and rotating the coordinates by a random offset to create random data bootstraps.

## Data, Materials, and Software Availability

Hybrid genomic data will be made publicly available upon publication. The previously published genomic data and the reference genome assembly and annotations were obtained from Nouhaud et al. (84)(https://doi.org/10.6084/m9.figshare.c.5332442.v1), and Nouhaud et al. (59) ENA project PRJEB51899. All code available upon request. All code will be made publicly available upon publication.

## Acknowledgements

The research was supported by an Academy of Finland grant 328961 and 346805 to JK. KL was supported by a European Research Council starting grant (ModelGenomLand 757648) and a grant from the Engineering and Physical Sciences Research Council UK (EPSRC, EP/X022595/1). We would like to thank Li Juan and Claudia Bank for their help and feedback on the X(2) analysis. We would also like to thank Dominik R. Laetsch and Sam Ebdon for their assistance with the initial steps of the gIMble analysis. Additionally we thank the Finnish Center for Scientific Computing (CSC) for computing resources used in this project.

