## Supplementary Materials and Methods for "Genomic incompatibilities are persistent barriers when speciation happens with gene flow in Formica ants"

**This PDF file includes:**

Supporting Text

Figures S1 to S14

Tables S1 to S9

**Supporting Text**

**gIMble analysis and detection of long-term barrier genomic regions**

Data used in gIMble

We used demographically explicit genome scans to detect barriers to gene flow with gIMble v1.0.3 (Laetsch et al. 2023). In this analysis, we used previously published diploid worker ant genomes sequenced from both parental species (Portinha et al. 2022). Details on library preparation and variant calling can be found in Portinha et al. (2022, mean sequencing depths ranging from 20.3 to 25.8). We investigated barriers using Finnish samples, since they are ecologically and evolutionarily most related to the hybrid samples used in the second analysis. We used four *F. aquilonia* (CF14a_1w, CF4b_1w, CF8b_1w, Pus2_1w) and four *F. polyctena* (Att1_1w, Jar6_1w, Lok3_1w, Fis2_1w) genomes, each sampled from a different population in southern Finland (Portinha et al. 2022). *F. polyctena* samples were collected from a region of sympatry and are known to be admixed (Portinha et al. 2022).

Demographic modeling to detect genomic regions of reduced long-term gene flow

BAM files and an unfiltered VCF file of the samples were obtained from Portinha et al. (2022) and were filtered and preprocessed for subsequent gIMble analyses using “gIMble preprocess” with default settings except for minimum coverage (--min_qual 1, --snpgap 2, --min_depth 5, and --max_depth 2 × mean coverage of each BAM).

First, we identified a “global” background model of demography for *F. aquilonia* and *F. polyctena* by fitting models with and without long-term gene flow. Specifically, we fitted isolation with migration (IM) models with both gene flow directions (*m_eF. aqu_* _→_ *_F. pol_* and *m_eF. pol_* _→_ *_F. aqu_*), a strict divergence (DIV) model without gene flow (*m_e_* = 0), and models of only migration (MIG) without divergence with both gene flow directions (*m_eF. aqu_* _→_ *_F. pol_* and *m_eF. pol_* _→_ *_F. aqu_*, T = ∞). gIMble uses the joint distribution of mutation types in short blocks, the block-wise site frequency spectrum (bSFS). We summarised data by the bSFS for block length 256b with kmax values of 2. Analyses based on the bSFS assume no recombination within blocks and a constant mutation rate (μ) across them. We assumed μ = 3.5 × 10^−9^ per site per generation, based on estimates available for social insects (Liu et al. 2017). The estimates of *T* were converted into absolute time using t = T × 2N_e_ × g, where N_e_ = θ/(4μ) and g is generation time. We assumed 2.5 generations per year, following Portinha et al. (2022). Note that the assumptions about μ and g do not affect the analysis, but simply scale *T* and N_e_ estimates in a way that is compatible with previous calibrations. We limited our analyses to intergenic and intronic (non-coding) sequences, excluding repeats, to minimize the direct effects of selection. Furthermore, chromosome 3 (Scaffold 3) was excluded from the global background analysis. In *Formica* species, this chromosome is known as the social chromosome, which contains multiple inversions and genes controlling whether a colony is led by one (monogynous) or multiple (polygynous) queens (supergene, Brelsford et al. 2020). Due to the inversions on this chromosome, recombination is strongly reduced between alleles of monogynous and polygynous colonies, leading to the maintenance of ancestral polymorphisms across *Formica* species which might bias demographic estimates. The best-fit global model involved long-term post-divergence gene flow from *F. aquilonia* into *F. polyctena* (IM*_F. aqu → F. pol_*, forwards in time; Table S1), the estimated migration rate being 1.50 individuals per generation. The species were estimated to have diverged 605,000 years ago. The estimated effective population sizes for *F. aquilonia*, *F. polyctena* and their shared ancestral population were 104,000, 72,000 and 116,000, respectively.

Second, the best-fit global model was fitted “locally” to detect genomic regions of reduced migration, i.e. barriers to gene flow. We used the same data in the local analysis as in the global analysis, except this time we included chromosome 3 in the analysis since it may contain barriers to gene flow. For the local analysis, a fixed number of non-coding blocks (-w 2000 -s 400) were grouped into windows with a minimum span of 32 kb. Because we used only non-coding blocks, i.e. coding and repeat sequences were excluded from the analysis, the real window span was usually greater than 32 kb (median 41.5kb for sympatry, Fig. S1). We investigated variation in *N_e_*s (*F. aquilonia*, *F. polyctena*, ancestral population) and *m_e_* across the windows by searching parameter combinations in a 12 × 12 × 12 × 16 grid. The grid was centered approximately on the estimates under the best fit global model (Table S1). The estimate of *T* was obtained from the global analysis, and fixed in the local analysis, that is, we assumed that the time of species divergence is a global event that is shared across the genome. We measured the local support for barriers via Δ_B0_ as proposed in Laetsch et al. (2023), i.e. positive Δ_B0_ values support reduced gene flow. To quantify the uncertainty in the parameter estimates, we used a parametric bootstrap in msprime (Baumdicker et al. 2022) to estimate the false positive rates (FPR) for Δ_B0_ at 0.05. We did this by simulating 100 window-wise datasets under a null model of a fixed (across windows) *m_e_* at the background level. For these simulations, we assumed a constant recombination rate based on one crossover per meiosis per chromosome, meaning that each chromosome has an averaged map length of 50 cM. Finally, we merged overlapping barrier windows (Δ_B0_>0 and FPR≤0.05) and referred to these as barrier regions.

**Imbalanced recombinant haplotype frequency analysis to detect BDMIs**

Data used in imbalanced recombinant haplotype frequency analysis to detect incompatibilities

The imbalanced recombinant haplotype frequency analysis (Li et al. 2022) was run using whole genome sequences from hybrid sexuals (males) collected at eight time points between years 2004 and 2022. Each time point has 30-40 individuals sampled (see Table S4 for samples). All samples were collected in the field during the emergence of reproductive individuals from nests within the W lineage of the Långholmen hybrid population in Hankoniemi, Southern Finland. Live adult individuals (recently emerged adults or late stage pupa) were brought to the lab in plastic bags with nest material and then transferred to centrifuge tubes filled with ETAX 99,5% ethanol and kept at -4 °C until DNA extraction. Overall, 447 samples were collected and processed for whole genome sequencing and variant calling. Only the male samples are used in this work, and included the 286 haploid male *Formica aquilonia* × *F. polyctena*, which were used for the imbalanced recombinant haplotype frequency analysis.

DNA extraction, library preparation and sequencing

DNA extraction was performed with Qiagen DNeasy Blood & Tissue Kit (Cat. No. / ID: 69504). Prior to DNA extraction, the ant tissue samples stored in alcohol were dried using a paper towel and one half from each individual was ground in microcentrifuge tubes using liquid nitrogen and plastic pestles. The lysis step was performed overnight without the optional RNase treatment. Concentrations of the DNA extracts were quantified using the fluorescence-based Qubit DS DNA HS kit (Q32851) from ThermoFisher Scientific.

DNA libraries were prepared after sample randomization (to avoid batch effects in the temporal dataset) with 150 ng of DNA from each individual DNA extract using New England Biolabs NEBNext® Ultra™ II FS DNA Library Prep Kit for Illumina (E7805L) according to manufacturer's instructions. Indexing of the samples was achieved using NEBNext® Multiplex Oligos for Illumina® Dual Index Primers Sets 1 and 2 (E7600S and E7780S, respectively). Manufacturer’s protocol for use with inputs ≥100 ng DNA was followed and DNA fragmentation was performed with the supplied enzymatic reagent for 11 minutes at 37°C and we aimed for a fragment size range of 200-450 bp; the recommended 15 minute incubation turned out to be too long. AMPure XP SPRI Reagent (A63881) from Beckman Coulter Life Sciences was used for the steps requiring magnetic beads and Invitrogen magnetic stand-96 (AM10027) from ThermoFisher Scientific was used to precipitate the magnetic beads. We aimed for the final DNA library size range of approximately 320-470 bp. Concentrations of the prepared DNA libraries were quantified similarly as the DNA extracts. Quality of the DNA libraries and their average sizes were analyzed with Agilent 5200 Fragment Analyzer and ProSize data analysis software.

Once the quality and the average size of each library passed QC, pre-pools were prepared from individual DNA library samples, with no more than 23 individual samples per pre-pool. Individual samples for these pre-pools were selected based on their average library size from fragment analysis, starting from the shortest and finishing with the longest average fragment size, so that each pre-pool contained samples of similar size range. These pre-pool samples were shipped on dry ice to Novogene Corporation Inc. for Illumina sequencing with the Illumina NovaSeq. We aimed for 2 GB of data for haploid males and 4 GB for diploid females.

Read mapping and variant calling for haploid males in the imbalanced recombinant haplotype frequency analysis

Raw 150 bp paired-end Illumina reads were trimmed with trimmomatic (v0.39; Bolger et al. 2014) with a minimum length of 80 and a leading and trailing minimum quality of 20. We used the annotated reference genome for a hybrid *F. aquilonia* x *F. polyctena* haploid male from Nouhaud et al. (2021). Mapping to the reference genome was run with bwa mem (v0.7.17; Li 2013, <https://github.com/lh3/bwa>) with default parameters. Overlapping reads were clipped with BamUtil(v1.0.15; Jun et al. 2015, <https://github.com/statgen/bamUtil>). Reads were quality checked with fastQC and visualized/summarized with multiQC (Ewels et al. 2016). Mean duplication and mapping rates across all samples were 13% and 97.6%. Mean coverages for male and female samples were 4.31× and 9.77× respectively.

Data for 447 individuals (142 females, 305 males) were used for SNP calling with FreeBayes v1.3.6 (Garrison and Marth 2012; <https://github.com/freebayes/freebayes>) with --limit-coverage 50, -n 4 (limit to 4 best alleles), and --no-population-priors. We normalized reads with vt normalize (Tan et al. 2015) and decomposed variants using vcfuniq from vcflib v1.0.3 (Garrison et al. 2022; <https://github.com/vcflib/vcflib>). We filtered variants to include biallelic SNPs with a minimum quality of 30, minimum SNP gap of 2, and balanced reads using bcftools (v 1.16; Danecek et al. 2021). Mean depth per individual was calculated with vcftools --depth (v0.1.17; Danecek et al. 2011) and then individuals were filtered to a minimum depth equal to half the mean individual coverage and a max depth equal to two times the mean individual coverage with bcftools (mean male coverage = 4.31×, mean female coverage = 9.77×). Sites in haploid male individuals were filtered to a minimum depth of 3 and female individuals filtered to a minimum depth of 6. Only sites typed in at least 60% of samples were retained for subsequent analyses. It is possible that morphologically male sexuals have diploid genomes when the sex determining region of a fertilized egg is homozygous. To identify if our samples contained diploid males, we counted the number of heterozygous sites in each male sample. Altogether we identified 19 male samples as diploids and dropped these samples along with female samples from further analysis, resulting in 2,736,443 SNPs in 286 haploid males.

Identifying and selecting Ancestry Informative Markers (AIMs)

The imbalanced haplotype frequency analysis described by Li et al. (2022) requires identifying SNPs where ancestry can be determined. We filtered the haploid male vcf file with bcftools to remove sites monomorphic for reference or alternate alleles and then converted the vcf file to .geno format with parseVCF.py (<https://github.com/simonhmartin/genomics_general>). AIM positions were identified using previously published data from Portinha et al. (2022): 10 female *F. aquilonia* (Sample IDs: CBAQ1_1w, CBAQ3_1w, CBAQ2_2w, Lai_1w, Lai_2w, Loa_1w, CF14a_1w, CF4b_1w, CF8b_1w, Pus2_1w) and 10 female *F. polyctena* samples (Sample IDs: CBCH1, CBCH2, CBCH3, CAGa, NAZa, VDa, Att1, Jar6, Lok3) from Europe . As in Li et al. (2022) we defined ancestry informative markers (AIMs) as SNPs displaying an allele frequency difference of 95% between the parental species. We filtered the haploid male .geno file to only include position identified as AIMs, and used a custom R script to convert genotype calls (A/A, T/T, C/C, G/G) to parental ancestry (0 for *F. polyctena* and 1 for *F. aquilonia*) and output .map and .ped files (<https://www.cog-genomics.org/plink/1.9/formats>) for input to the AD scan analysis. This resulted in a total of 21,283 AIMs across the genome (Fig. S7).

Detecting BMDIs in a natural hybrid population

Detection of incompatibilities (BMDIs) is based on an imbalance of recombinant haplotype frequencies in hybrid genomes, searching for heterospecific combinations of SNPs where *g_10_*>*g_11_*, *g_00_*>*g_01_* or *g_10_*>*g_11_*, *g_00_*>*g_01_* where *g* is the frequency of ancestral allele combinations between parent 1 and parent 0. This method has good resolution to detect negative epistasis between loci in relatively young hybrid populations with equal admixture proportions. The Långholmen hybrid population in southern Finland is thus ideal, as it is 24-50 generations old and has roughly equal admixture proportions (ca. 45% contribution from *F. aquilonia*, (Nouhaud et al. 2022b). The analysis was run using the ancestry informative markers (AIMs) identified as above in haploid males, removing the need for phasing. We calculated *X*(2) and D’ statistics (Li et al. 2022) for all pairwise AIM combinations using the fishStat.permutation R script from Li et al. (2022) ([zenodo.org/doi/10.5281/zenodo.6334595](http://zenodo.org/doi/10.5281/zenodo.6334595)). To identify regions of the genome which are putative BDMI pairs, we performed a sliding window analysis which compared the focal window to each non-overlapping window across the genome. As the identified AIMs were not evenly distributed across the genome (Fig. S7), windows used a set number of consecutive AIMs to prevent AIM density affecting inferred incompatibilities. We selected 20 AIMs per window as this was the closest average window size we could get to the observed LD decay of 50Kb while maintaining a feasible number of comparisons. While there was a large variance in window sizes (Table S5), There was not a strong correlation between window size and X(2) (Fig. 14).

For each window pair we calculated mean *X*(2) and mean D’ between the AIMs in each window, as well as the fraction of AIM comparisons with X(2) < -0.005 (Fig. S8B). Given the number of comparisons between window pairs (window size^2^), we performed 1000 resamples of an equal number of X(2) values from the complete set of AIM comparisons across the genome, and calculated the fraction of AIM comparisons with X(2) < -0.005 to create a null distribution of how many candidate AIM pairs are expected between two windows by chance. We also performed 1000 random resamples of 40 AIMs across the genome and calculated the number of sites with at least one comparison with X(2) < -0.005.

We considered a window pair to be a putative BDMI if it met the following criteria:

1. The average X(2) between the two windows is < -0.005.
2. The fraction of AIM comparisons between the two windows is greater than the 99th percentile of the bootstrapped fraction distribution.
3. The number of total candidate AIM positions in the two windows is greater than the 99th percentile of the bootstrapped distribution.

These criteria assume the true BDMI loci will distort alternative recombinant haplotype frequencies not only at the incompatible locus, but due to linkage create a similar signal around the BDMI loci (Li et al. 2022; Szabo and Cutter 2024). By averaging the X(2) signal across larger regions we aim to distinguish signal between BDMIs and spurious associations, as spurious associations are unlikely to have linked signals and will average out with combined X(2). Meanwhile around true BDMIs a negative X(2) value will be maintained when combining comparisons between groups of linked SNPs. With this method we expect to capture loci as candidates which are linked to true incompatibility loci, but not necessarily causal SNPs themselves, and interpret candidate BDMI regions identified as those impacted by a BDMI present within them.

Quantifying incompatibilities

We quantified the number of incompatibilities simply as the total number of BDMI pairs meeting the BDMI candidate criteria. Based on simulations in Li et al. (2022), the -0.005 cutoff for X(2) distinguished neutral and BDMI loci. However, to understand how more filtering affects our results, we present the number of candidate BDMIs pairs for 23 cutoff values (-0.0600, -0.0575, -0.0550, -0.0525, -0.0500, -0.0475, -0.0450, -0.0425, -0.0400, -0.0375, -0.0350, -0.0325, -0.0300, -0.0275, -0.0250, -0.0225, -0.0200, -0.0175, -0.0150, -0.0125, -0.0100, -0.0075, -0.0050), where each cutoff contains the BDMI pairs where the average X(2) between windows is less than or equal to the cutoff.

Calculating multi-locus interactions and network features

To calculate the regression of BDMI degree and *m_e_* we calculated the number of BDMI candidates at each locus with bedtools merge using the -c = 1 and -o = “count” to count the number of BDMI regions at each locus in the BDMI bed file. We then calculated *m_e_* for each barrier region with bedtools intersecting the BDMI and gIMble bed files with -wa and -wb. We calculated the correlation between BDMI and *m_e_* using the cor.test function from the stats R library (R Core Team 2025). We repeated this procedure for each of the 23 X(2) thresholds.

To check for significance of the observed slopes we bootstrapped p values by randomizing *m_e_* values across the observed BDMI degrees and recalculating the correlation coefficient. We repeated this procedure 1000 times and compared the observed slope to the bootstrapped distributions of slopes using the ecdf function from the stats R library (R Core Team 2025) to calculate p values. When the observed value was greater or less than any bootstrap p was determined to be <1/1000 resamples, or 0.001.

To calculate the network measures (Fig. 3E-H, Fig. S13) for the BDMI network at each X(2) threshold we used the igraph library in R (Csárdi et al. 2025; R Core Team 2025). We calculated the degree distribution of each network with the function degree and took the max value to be max degree. We calculated the average path with the mean_distance function and we calculated the average betweenness with the betweenness function.

To compare the observed network to a relevant BDMI model for speciation we chose the Orr (1995) extension of the model. To translate this to a network, we started with one node, equivalent to one locus where a substitution has occurred between two otherwise identical lineages. Nodes were added sequentially, and interacted with previous nodes at rate *p*. We determined *p* as the mean node degree (eq.1) of the BDMI network at each X(2) threshold where d bar is the mean degree, and |*V*| is the number of vertices in the graph.

$p = \frac{\underline{d}}{\left| V \right|-1}$	(1)

As *p* is stochastic, we generated 1000 Orr networks to generate a null distribution of network measures to compare the BDMI network against. We note that the observed *p* is far higher than the estimates of the probability of interactions between loci as noted in Orr and Turelli (2001). However, our goal is to compare the network topology generated by the model, rather than the number of incompatibilities.

**Overlap between long-term barriers detected with gIMble and BDMIs identified by imbalanced haplotype frequency analysis**

To test for an enrichment in overlap between gIMble barriers and candidate BDMI regions (Fig. 1C) we performed a circularization bootstrap similar to Yassin et al. (2016), Nouhaud et al. (2022), and Ebdon et al. (2024) using a custom R script and bedtools v2.31.0 (Quinlan and Hall 2010). Both gIMble and the AD scan analyses used rely on SNP data, meaning that the first data point for each analysis does not start at the same position as the start of chromosome. To remove bias in overlap caused by functional features (e.g. CDS) outside of the range of data available for each analysis, we trimmed all input files to the overlap script to the innermost start and end SNP of either the gIMble or AD scan data as appropriate. After this procedure we simply used the new start and end as the chromosome length for bootstrapping. This procedure should not impact the results much, as >98% of each chromosome length remained.

For bootstrapping, features from each input bed file are merged with bedtools merge (merges overlaps within files), and then the total observed overlap between the two files is calculated in base pairs with bedtools intersect. We then circularize each chromosome, shift the features of the first input bed file by a random percent of each chromosome length, and recalculate the overlap with bedtools intersect. Repeating the circularization 1000 times creates a bootstrapped null distribution of total overlap values while preserving the distances/clustering of each input feature within chromosomes. A p-value for each observed value versus the bootstrapped distribution is calculated with the ecdf function from the stats R library.

Annotation Features

We used the same bootstrapping procedure as above to calculate the overlap between gIMble barriers and gene and repeat annotations from Nouhaud et al. (Nouhaud et al. 2022a)(2021) ([doi.org/10.6084/m9.figshare.c.5332442.v1](http://doi.org/10.6084/m9.figshare.c.5332442.v1)). We filtered the gene annotations file to only include CDS regions. We calculated intronic regions as gene annotation regions - CDS annotation regions (i.e. we assumed all gene regions were either CDS or intron). The annotation of repeat regions was used as is, and includes multiple types of repeats, including simple repeats, satellite DNA, and transposable elements (TEs). We repeated this procedure for each BDMI X(2) cutoff.

We divided this analysis into two sets. One set including overlapping genome-wide gIMble barriers and BMDIs with CDS, introns, and repeats, and a second set including only regions where gIMble barriers and candidate BDMIs overlapped (long-term BMDIs). To calculate this second set we created a bed file by intersecting the gIMBle bed file with each BMDI X(2) cutoff using bedtools intersect (Quinlan and Hall 2010), and bootstrapping the overlap as above.

Identifying centromeres

To characterize the positions of centromeres, we mapped a 129bp satellite DNA marker, found in the centromere regions in *Formica* ants (Lorite et al. 2004), against the reference genome using BLASTn v2.16.0+; Camacho et al. 2009). If the satellite DNA marker was not found on a chromosome, we identified the centromere regions in the chromosome level assembly of a related species, *F. rufa* (iyForRufa2.1), and then used this information to define the likely chromosome end the centromere belongs to by identifying conserved synteny between the species using RagTag (Alonge et al. 2022).

As a proxy for recombination, we tested whether BDMI and gIMble barriers are closer to centromeric regions than expected by chance. To do this, we calculated the average central position for each centromere for each scaffold which had data. We then took the absolute value of the difference of the center of each BDMI and gimble region to the centromere. To test if this distribution of distances was smaller than expected by chance, we circularized per chromosome and shifted coordinates by a random percent of each chromosome length and recalculated the distance to centromeres. We compared the observed distribution to the distribution of 1000 circularized resamples with the Wilcox rank sum non-parametric test. We implemented this with the wilcox.test function in R and the alternative hypothesis “lesser” to test if the observed median is significantly lower than the bootstrapped median.

Hybrid index and barrier sorting

An important aspect of hybrid genome evolution is how the mixed genomic ancestry of an initial hybridization event sorts over time in a population. We expect allele combinations to emerge which are compatible. As alleles of shared ancestry have been previously tested together in parent genomes, regions of single ancestry across the population begin to emerge (Nouhaud et al. 2022b). To understand how ancestry sorted in the 286 hybrid males we calculated mean *D̄* across barrier and non barrier windows. Using the same 10 *F. aquilonia* and 10 *F. polyctena* samples as in the AIM identification above, we filtered for sites with a difference in ancestry of <0.2 or >0.8 resulting in 65,970 sites across the genome. To prevent an effect of window size, non-barrier windows (mean 1.6Mb) were split into multiple windows targeting a mean size equal to the mean of the true gIMble barrier windows (61Kb). We then calculated mean HI for each individual in each region and took the variance across individuals as *D̄* (Barton and Gale 1993). To summarize the data we took mean *D̄* across all barrier and non-barrier windows (Fig. 4C). We compared the observed mean *D̄_barrier_* to mean *D̄_non-barrier_* by bootstrapping null distributions for each and calculating p values using circular resampling as above.

Identifying candidate incompatibility genes

To identify candidate incompatibility genes we filtered the gIMble and X(2) results following two criteria. We extracted gene annotations from regions where:

1) gIMble Δ_b0_ > 0 and *m_e_* = 0 (i.e. regions acting as complete barriers to gene flow)
2) Regions where both ends are a candidate BDMI pair.

The intersect of the paired BDMI data and gIMble was run with bedtools pairtobed and type = “both”. This filtered to 13 gene annotations (Table S9). Annotations from the gff3 file were checked on both strands (-B), filtered for min length 30 (-l), we discarded mRNAs with inframe stop codons (-V), reading frame was shifted if the original transcript led to an inframe stop codon (-H), discarded mRNAs with non-canonical splice sites (-N), and removed mRNAs without both a START and terminal STOP codon (-J) gffread (Pertea and Rozenberg 2020). The filtered annotation file was converted to amino acid sequences with gffread -y.

To get gene descriptions we blasted the candidate amino acid sequences against the NCBI nucleotide database (nt) using tblastn (Camacho et al. 2009) (Table S9) and selected the top hits based on e-value.

**Supporting Figures**

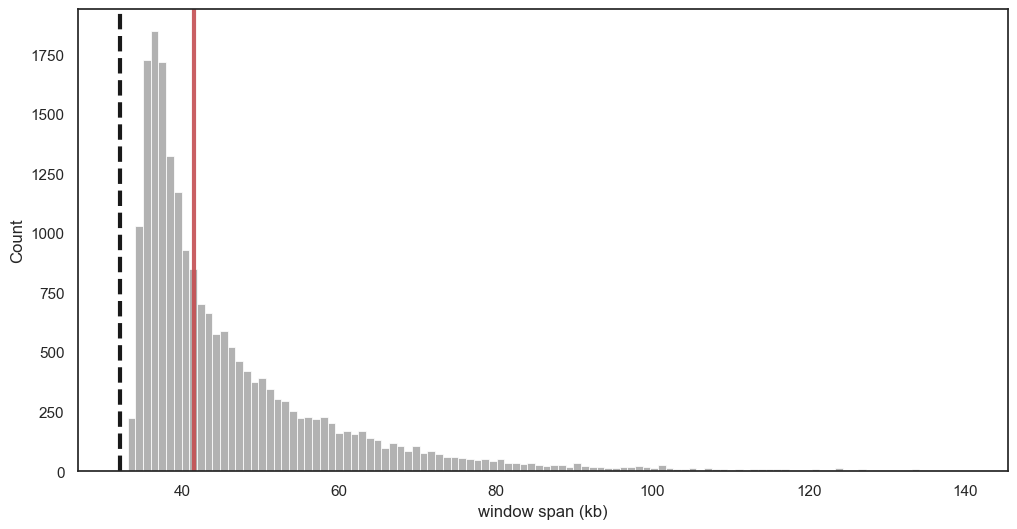

**Figure S1.** The window span distribution of the data set used in gIMble analyses. The genome was analyzed in sliding windows of non-coding sequence. We used a window span of 32 kb (dashed black line), but given that we used only non-coding sequences the window span is greater than that, varying from 33 kb to 756 kb. Median window span is shown with a solid red line (42 kb).

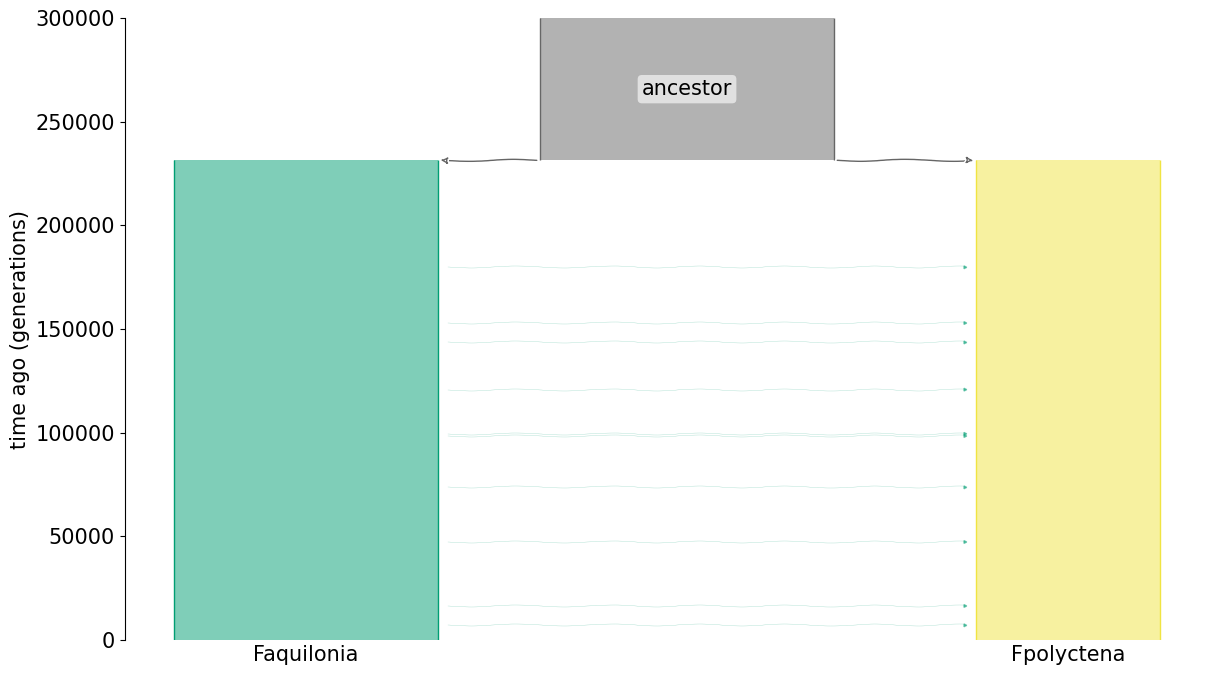

**Figure S2.** Illustration of the best fit demographic model between *F. aquilonia* and *F. polyctena*, IM*_F .aqu→F. pol_*. The block widths indicate relative effective population sizes (*N_e_*) of the ancestral population and its two descendants. Arrows indicate unidirectional gene flow at a rate of 1.5 migrants per generation from *F. aquilonia* to *F. polyctena*. The figure was produced with demesdraw (Gower et al. 2022).

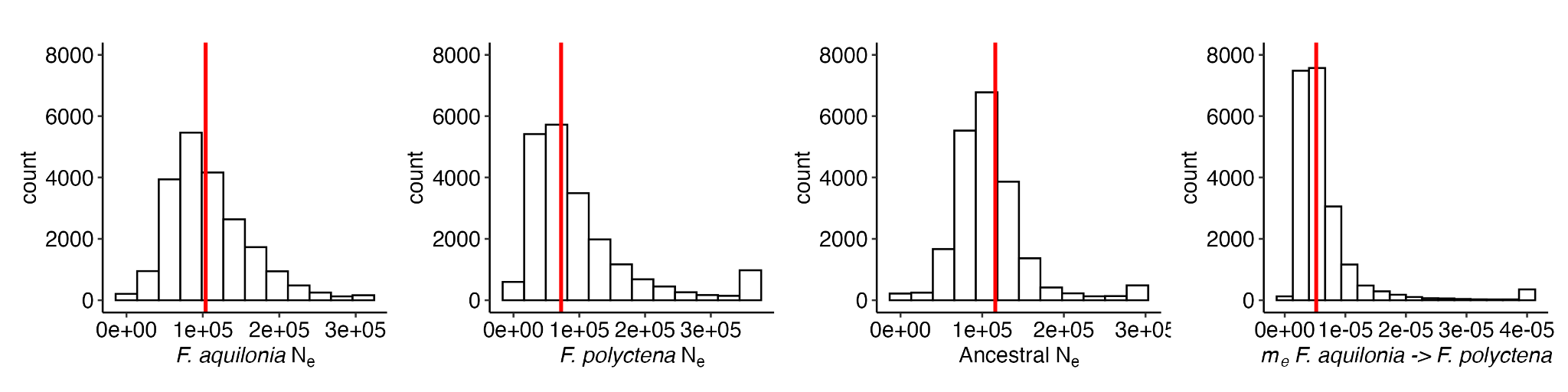

**Figure S3.** Variation in the effective population sizes (*Ne*_e_) of *F. aquilonia*, *F. polyctena* and their shared ancestral population, and variation in the migration rates (*m*_e_) from *F. aquilonia* to *F. polyctena* across sliding windows. The red vertical lines indicate estimates under the global model (Table S1). The number of bins in each distribution corresponds to points in the 12 × 12 × 12 × 16 parameter grid used for inference.

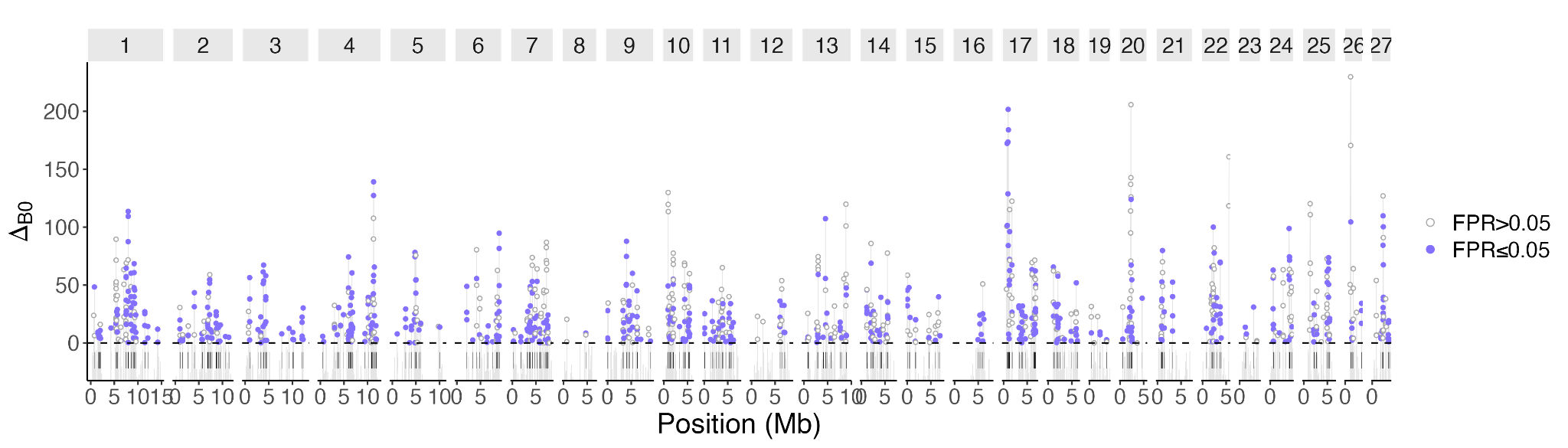

**Figure S4**. Long-term barriers to gene flow between *F. aquilonia* and *F. polyctena* identified by gIMble across all 27 chromosomes. Dots indicate windows with Δ_B0_ > 0, i.e. candidate barriers to gene flow, where a history of reduced *m_e_* fits better than a model assuming the global estimate. Closed dots indicate significant barriers (false positive rate, FPR ≤ 0.05), and open dots false positives (FPR > 0.05). Significant barrier regions (overlapping windows with Δ_B0_ > 0 and FPR ≤ 0.05) are marked with vertical bars.

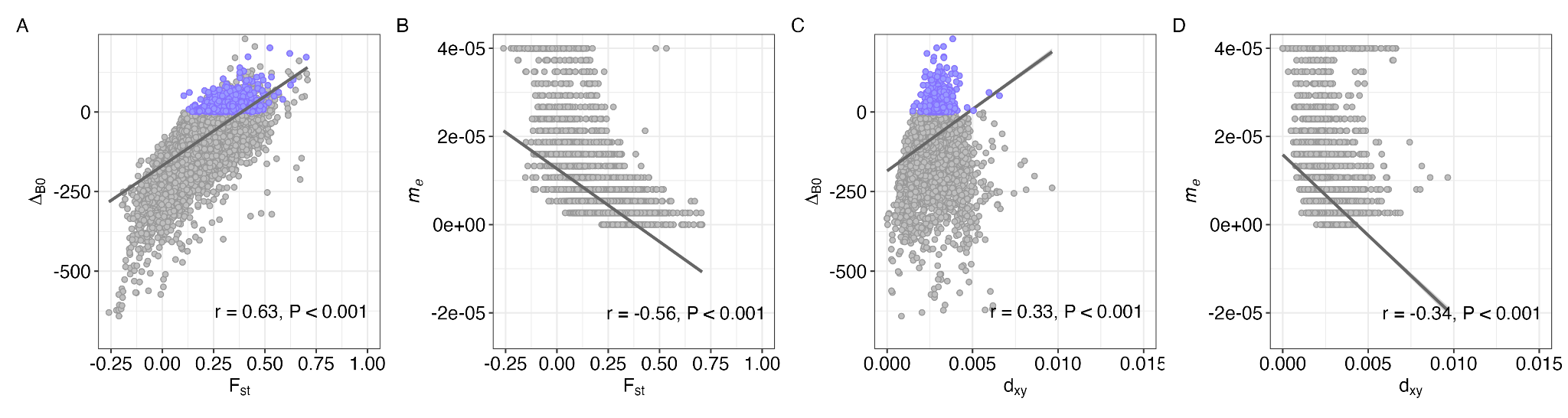

**Figure S5**. Window-wise Pearson correlations (r) between metrics of barrier loci (*m_e_*, Δ_B0_) and genetic divergence (F_st_ and d_xy_). (A) F_st_ and Δ_B0_, (B) F_st_ and *m_e_*, (C) d_xy_ and Δ_B0_, and (D) d_xy_ and *m_e_*. Closed dots indicate barriers to gene flow (Δ_B0_ > 0 and FPR ≤ 0.05). Lines represent regression lines.

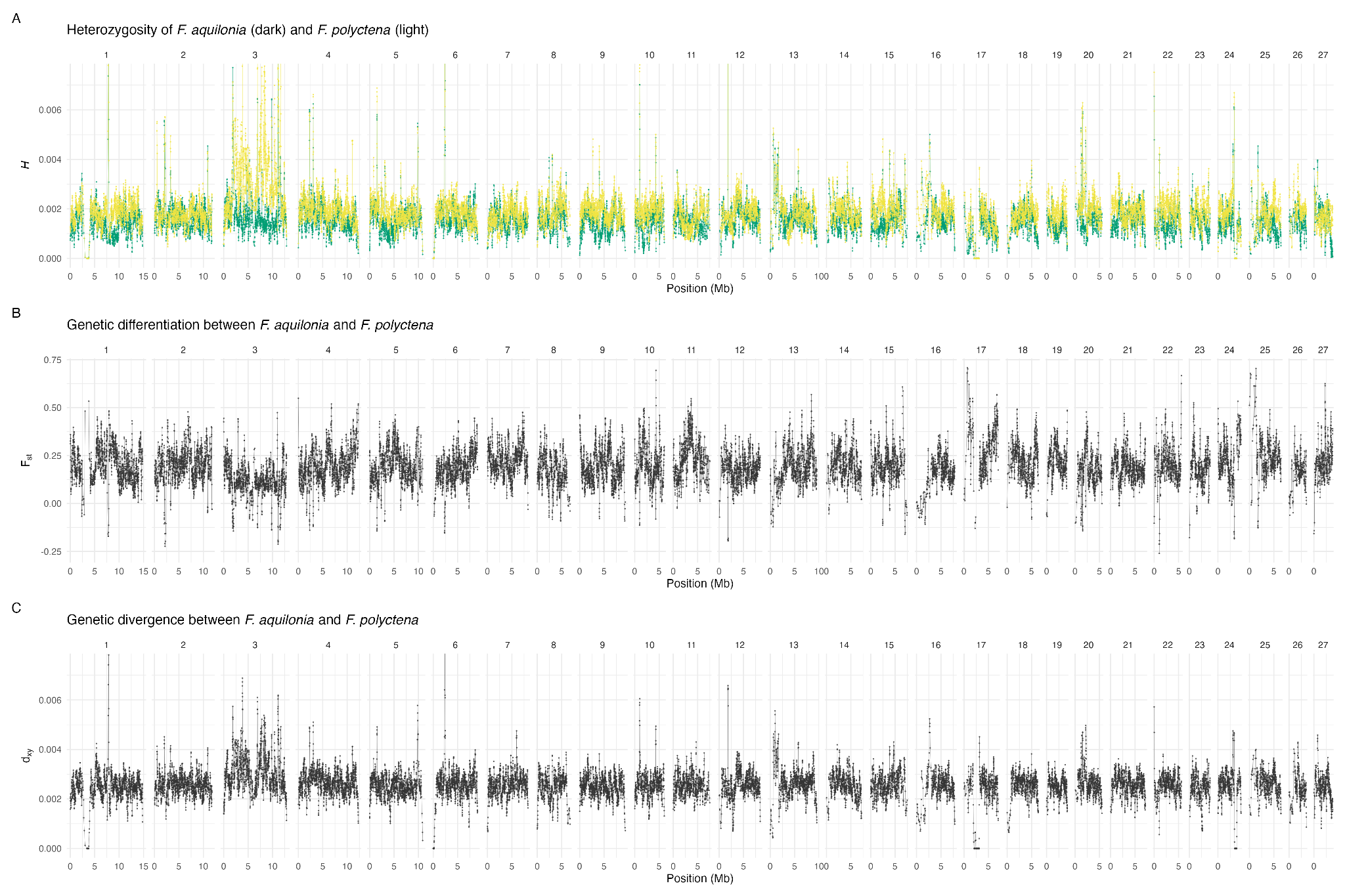

**Figure S6.** Genome-wide (A) heterozygosity (*H*) of *F. aquilonia* and *F. polyctena*, and (B) genetic divergence (d_xy_) and (C) genetic differentiation (F_st_) between the species.

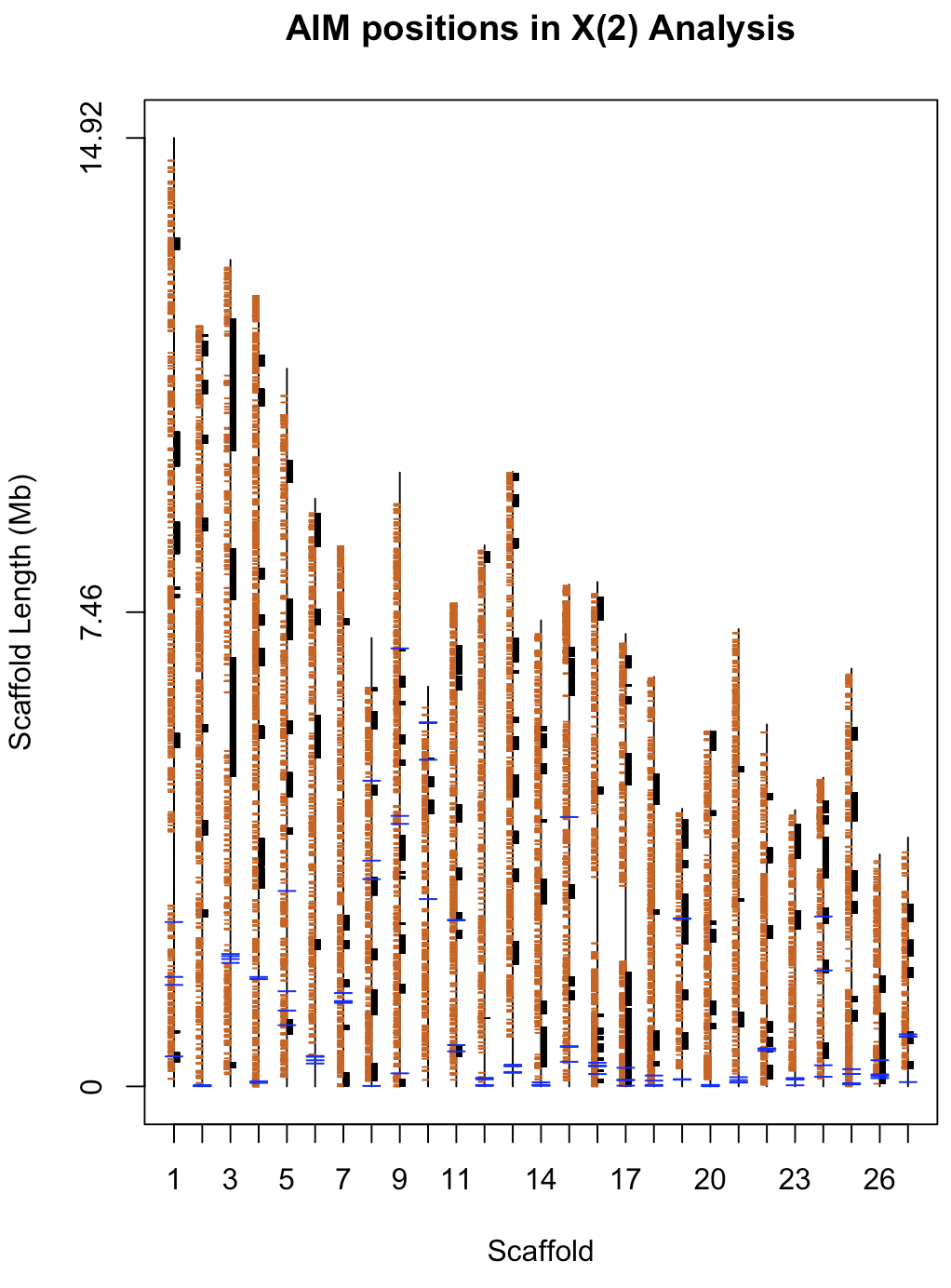

**Figure S7**. Map of 21,283 AIMs used in the unbalanced recombinant haplotype frequency analysis. Black vertical lines are scaffolds, small orange horizontal dashes are AIM positions, larger blue horizontal dashes are reference assembly gaps, black rectangles are candidate BDMI regions at the X(2) < -0.035 threshold.

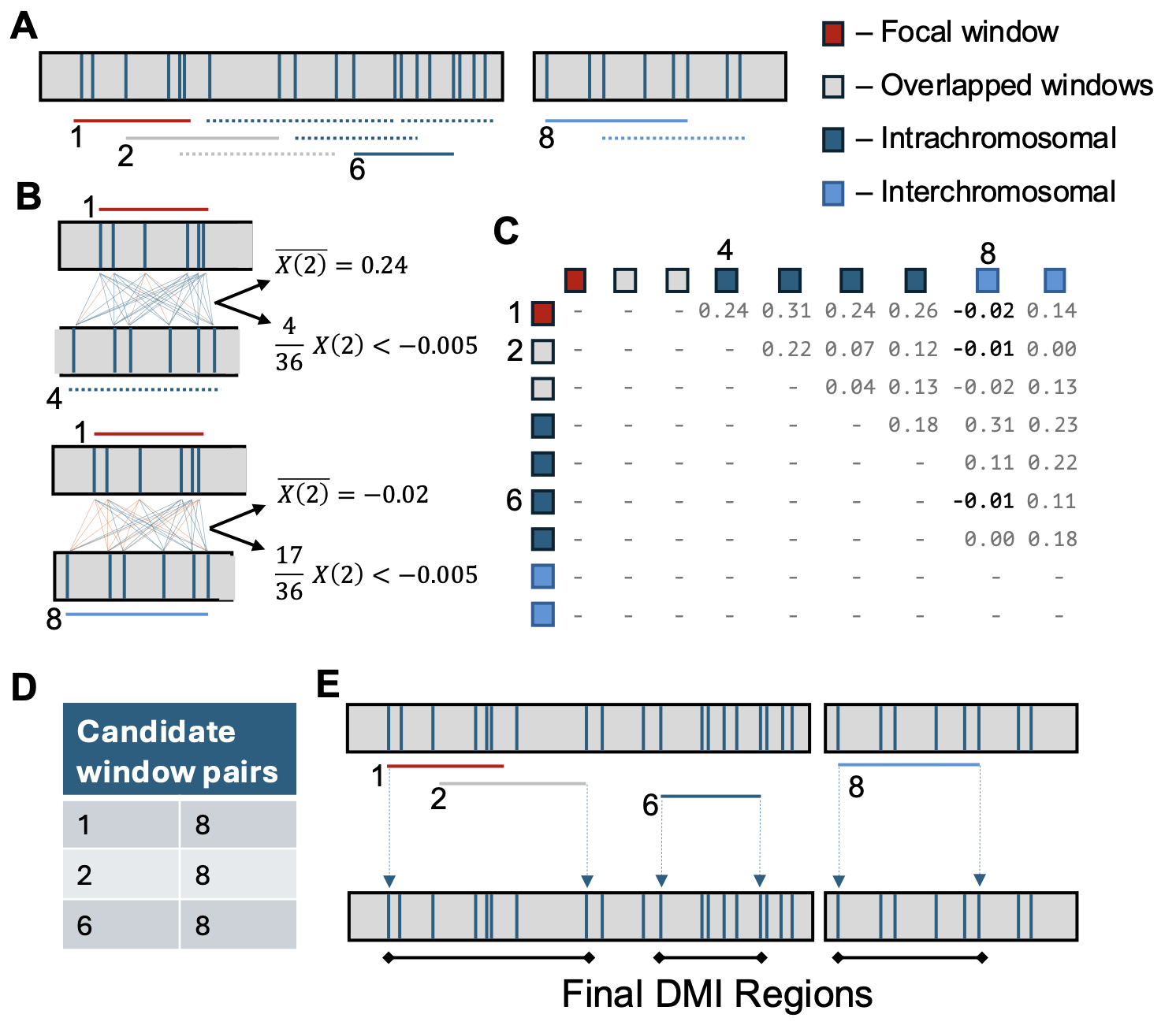

**Figure S8.** Identifying candidate BDMI Regions. (A) Windows contain equal numbers of AIMs and overlap by a step size of 5% (e.g. windows of 20 AIMs shift in steps of 4, with the last last window containing 18-22 AIMs (20 ± step size / 2) ). (B) Example comparison between focal window 1 (red line) and an intrachromosomal window 4 (dark blue dash). The mean X(2) between all AIMs in each window is calculated, as well as the fraction of AIM comparisons with X(2) < -0.005 (Li et al. 2022). (C) Comparisons are pairwise between all non-overlapping windows (i.e. window 1 shares AIMs with windows 2 and 3, so no comparison is made). (D) Candidate window pairs are those with a signal of a BDMI as indicated by both negative mean X(2) and a fraction of comparisons which is greater than expected by chance (bolded values in (C)). (E) Candidate window pairs which share AIMs (e.g. pairs 1 & 8 and 2 & 8) are merged to create the candidate BDMI regions.

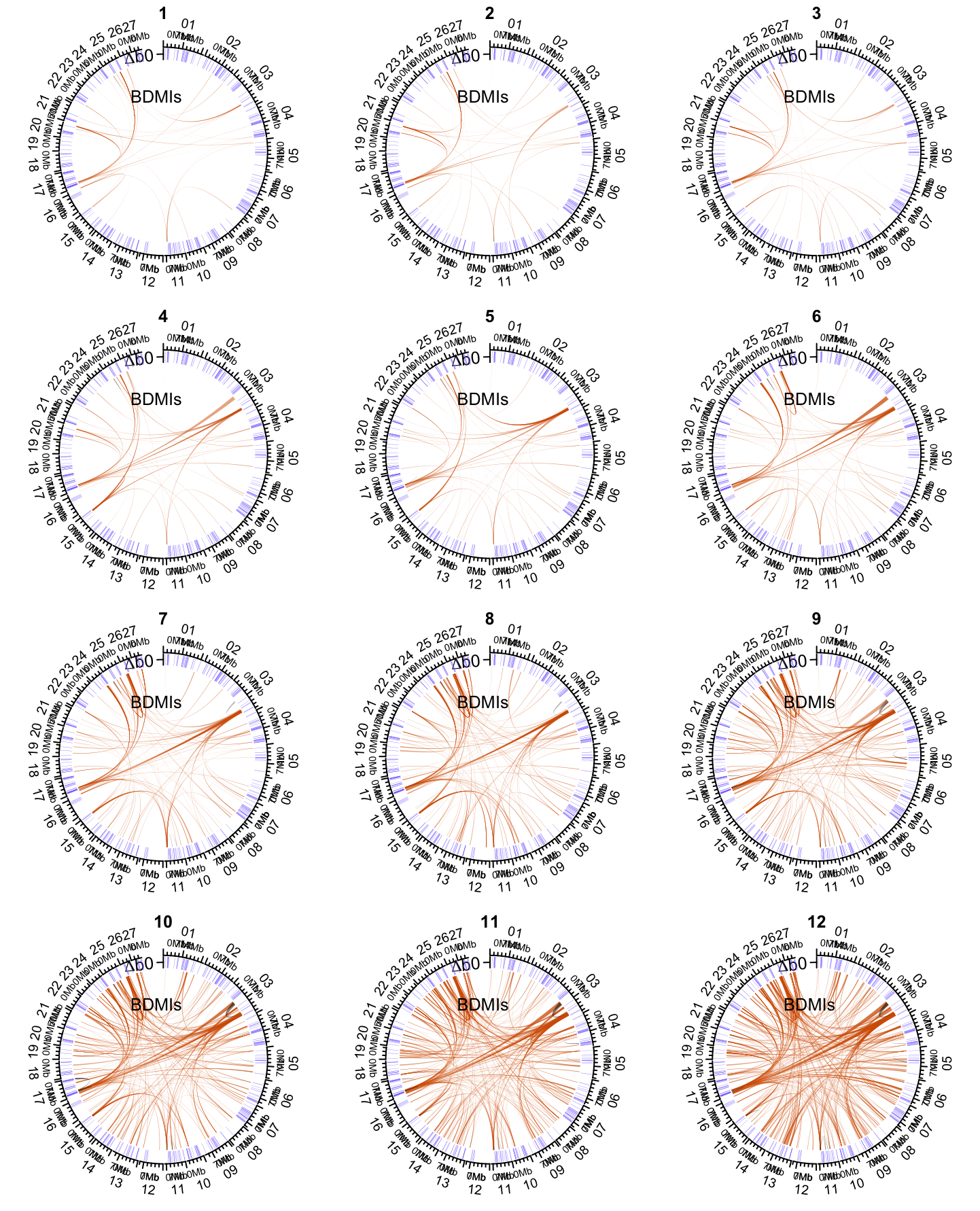

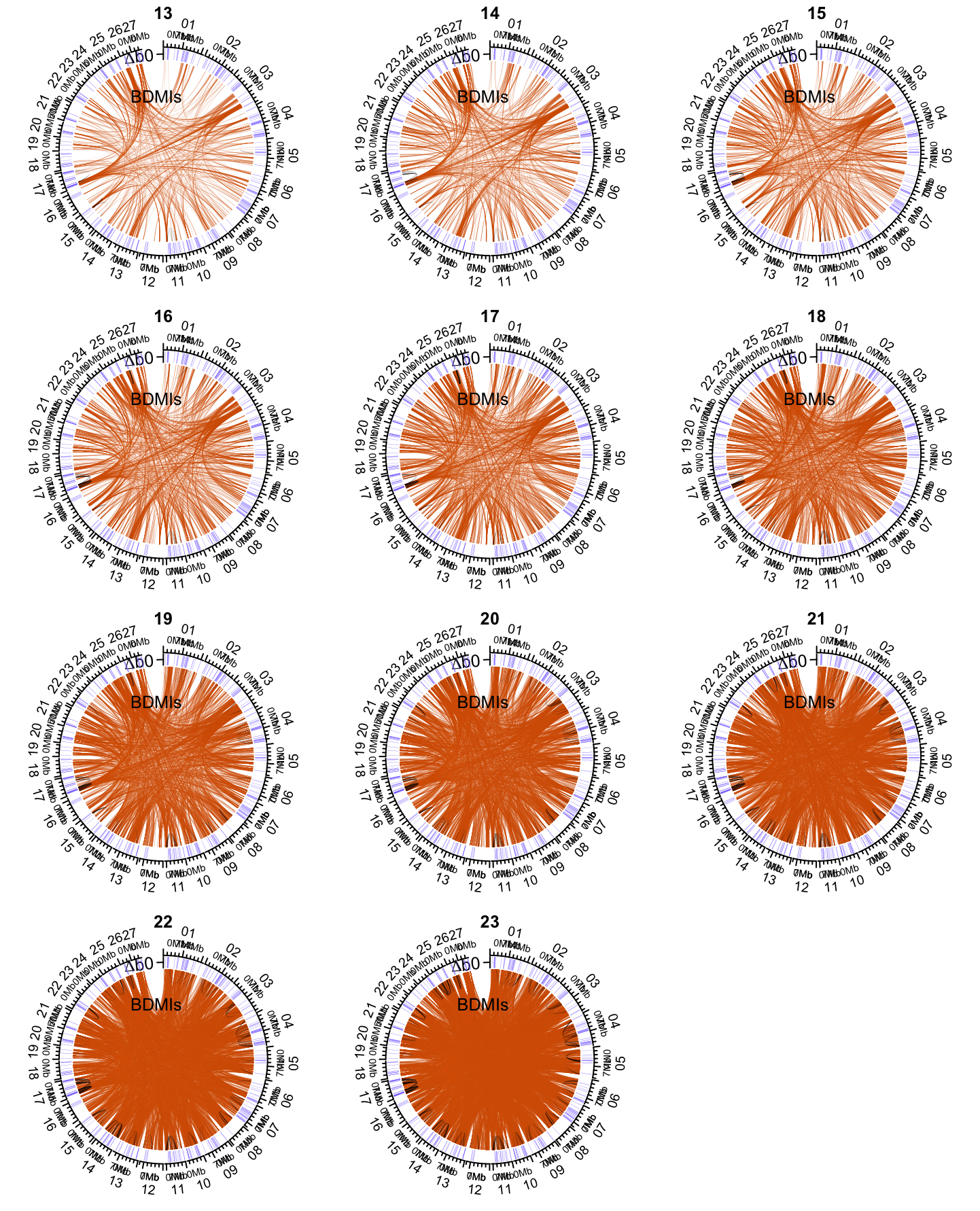

**Figure S9.** Circos plots for all 23 X(2) cutoff values used in the imbalanced haplotype frequency analysis.

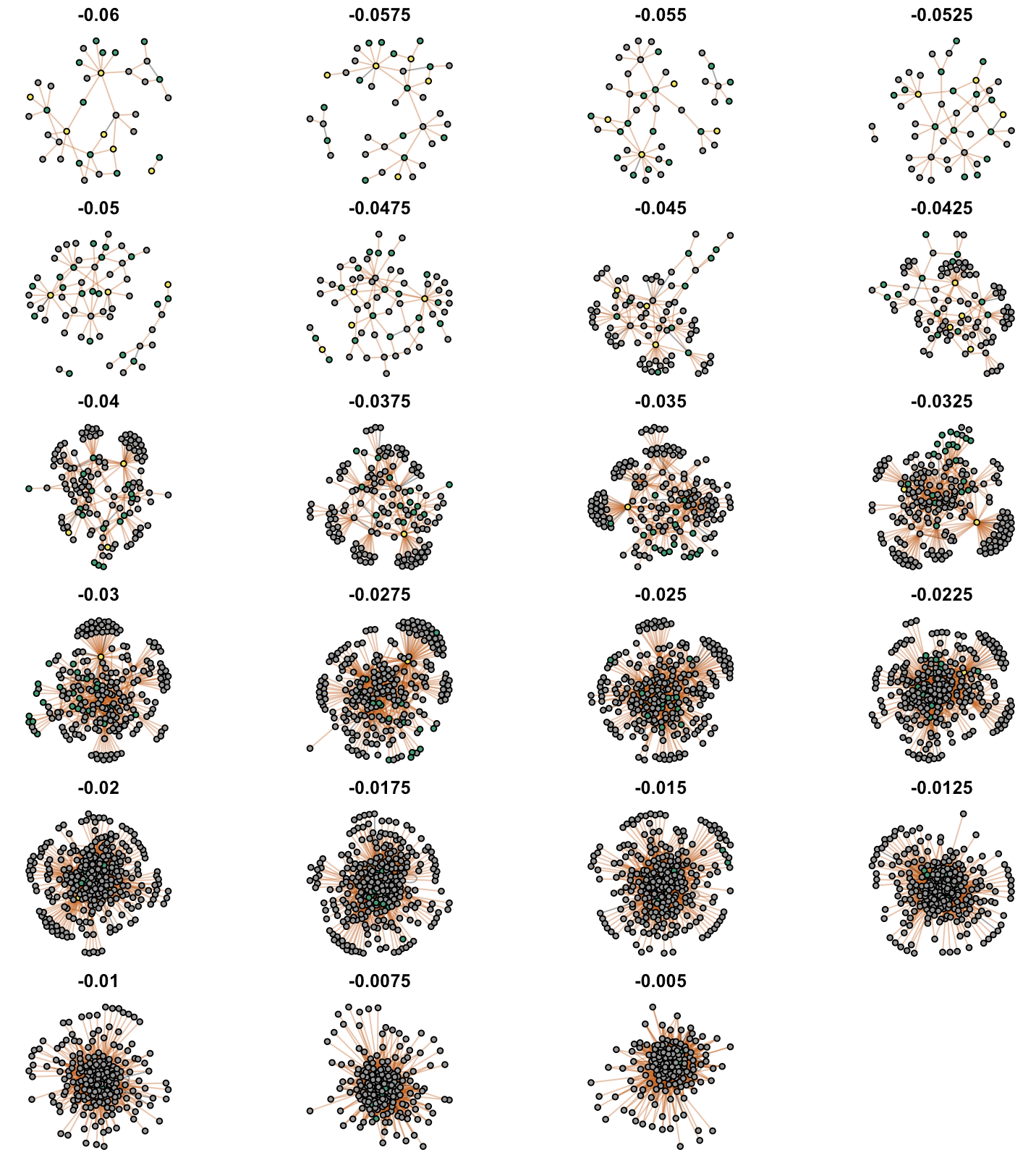

**Figure S10.** Network visualizations for all X(2) thresholds. Nodes are colored based on mean ancestry proportion within the BDMI region the node is associated with. Regions with a hybrid index of <=0.2 are colored green (primarily F. aquilonia ancestry), regions with a hybrid index >= 0.8 are colored yellow (primarily F. polyctena ancestry), regions with intermediated hybrid indices >0.2 & <0.8 are colored grey.

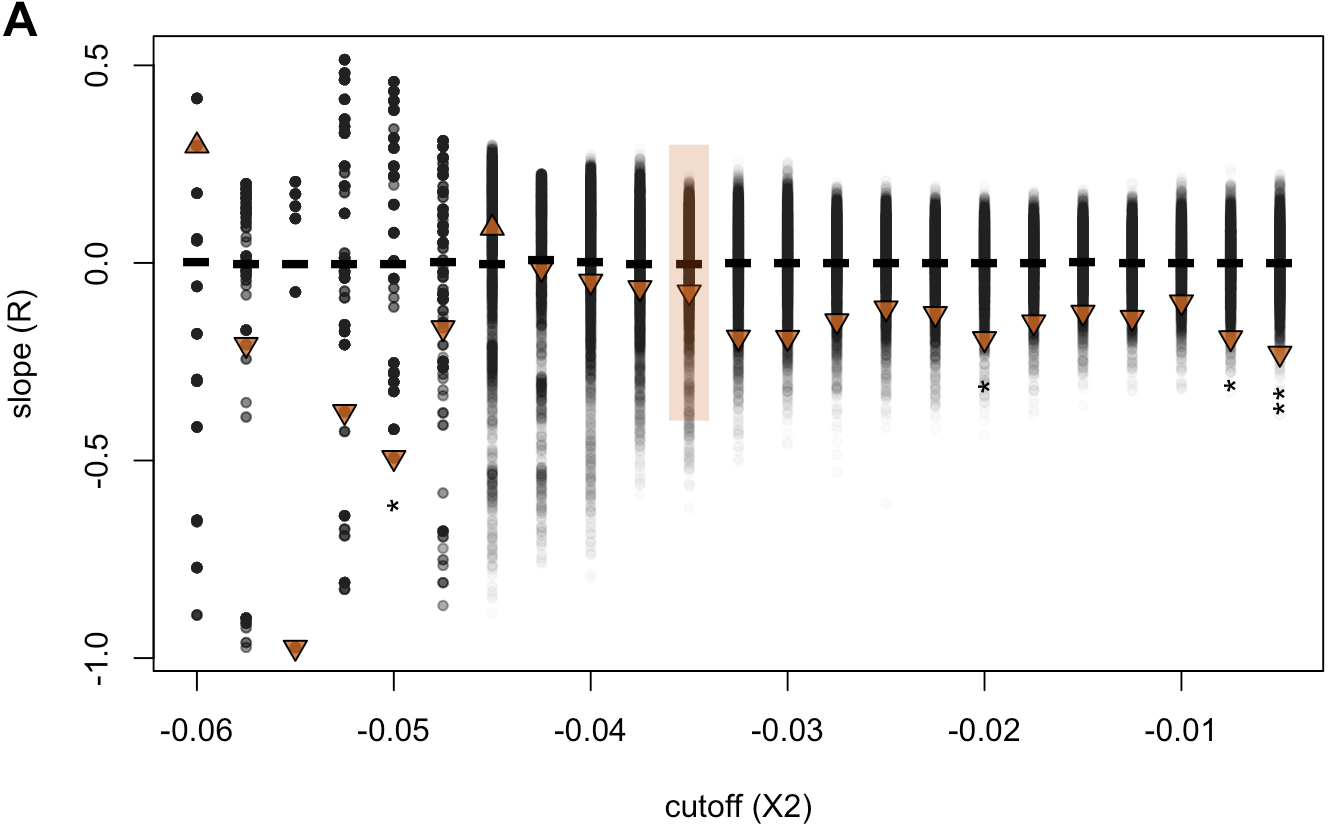

**Figure S11.** The effect of removing Scaffold 17 on the correlation between BDMI degree and long-term barrier strength (*m_e_*). Negative correlation remains for X(2) > -0.0425 and significant for X(2) = -0.05, -0.02, -0.0075, and -0.005.

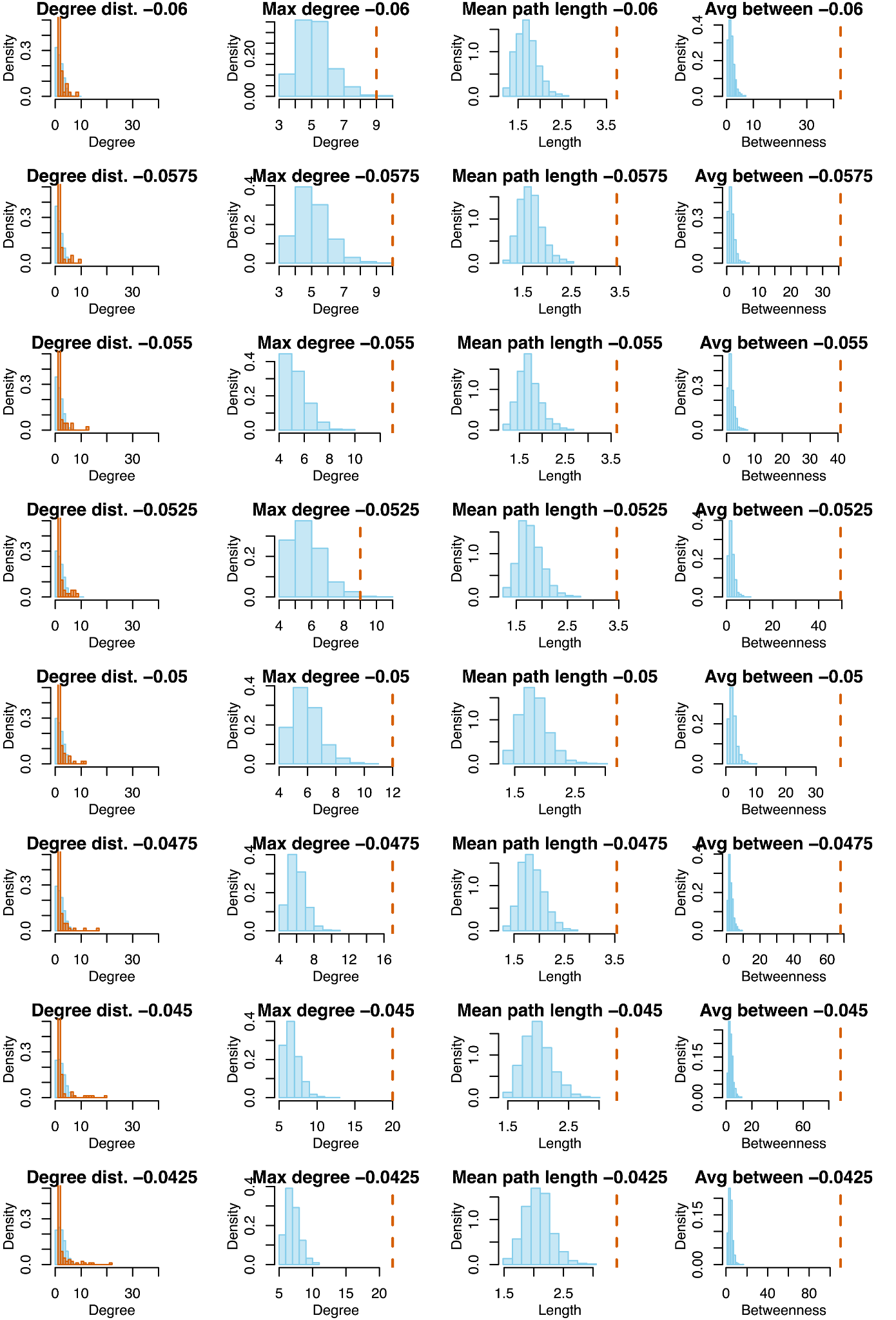

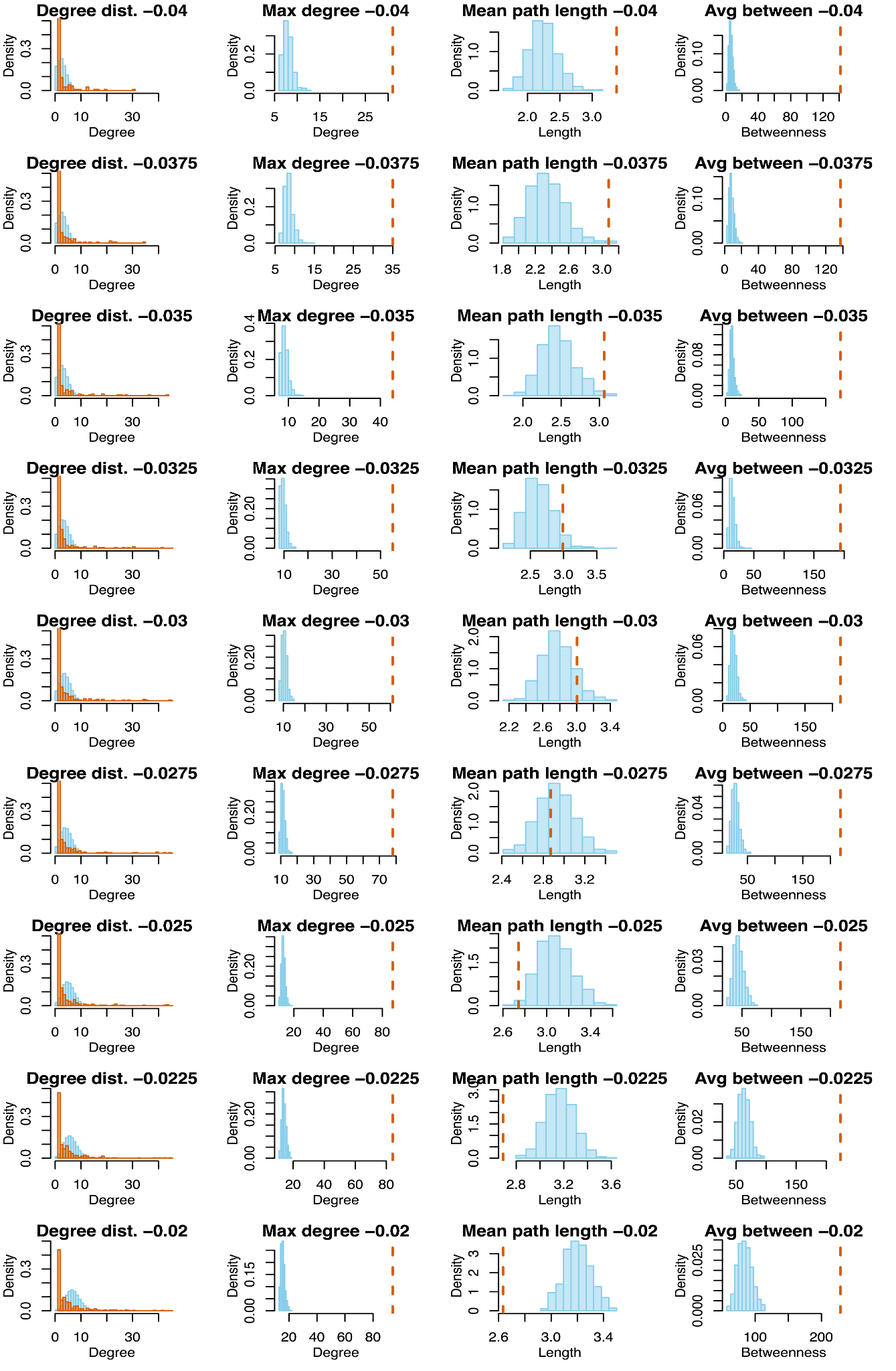

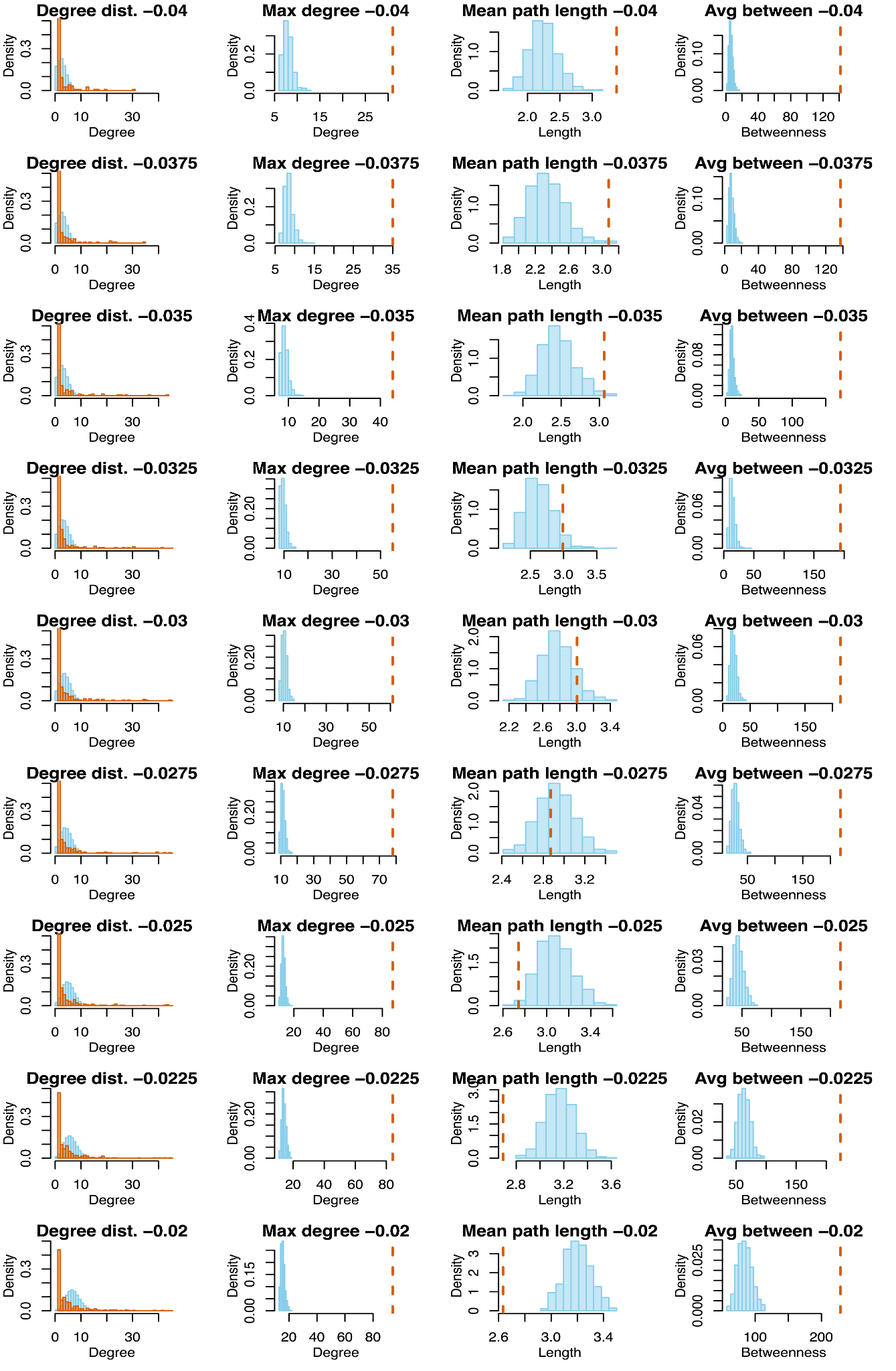

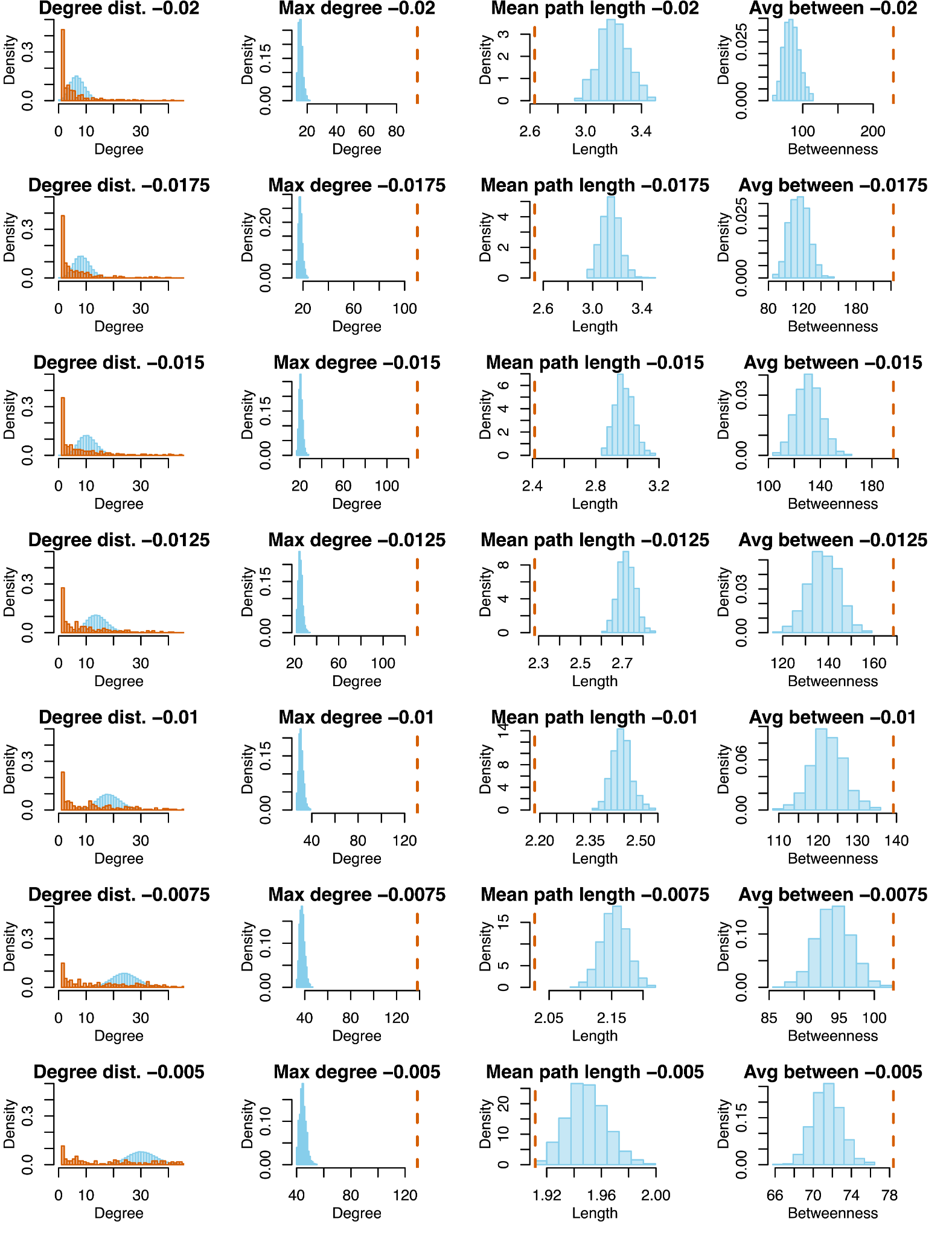

**Figure S12.** Network measure visualizations for all X(2) thresholds. See main text and Fig. 3 for description.

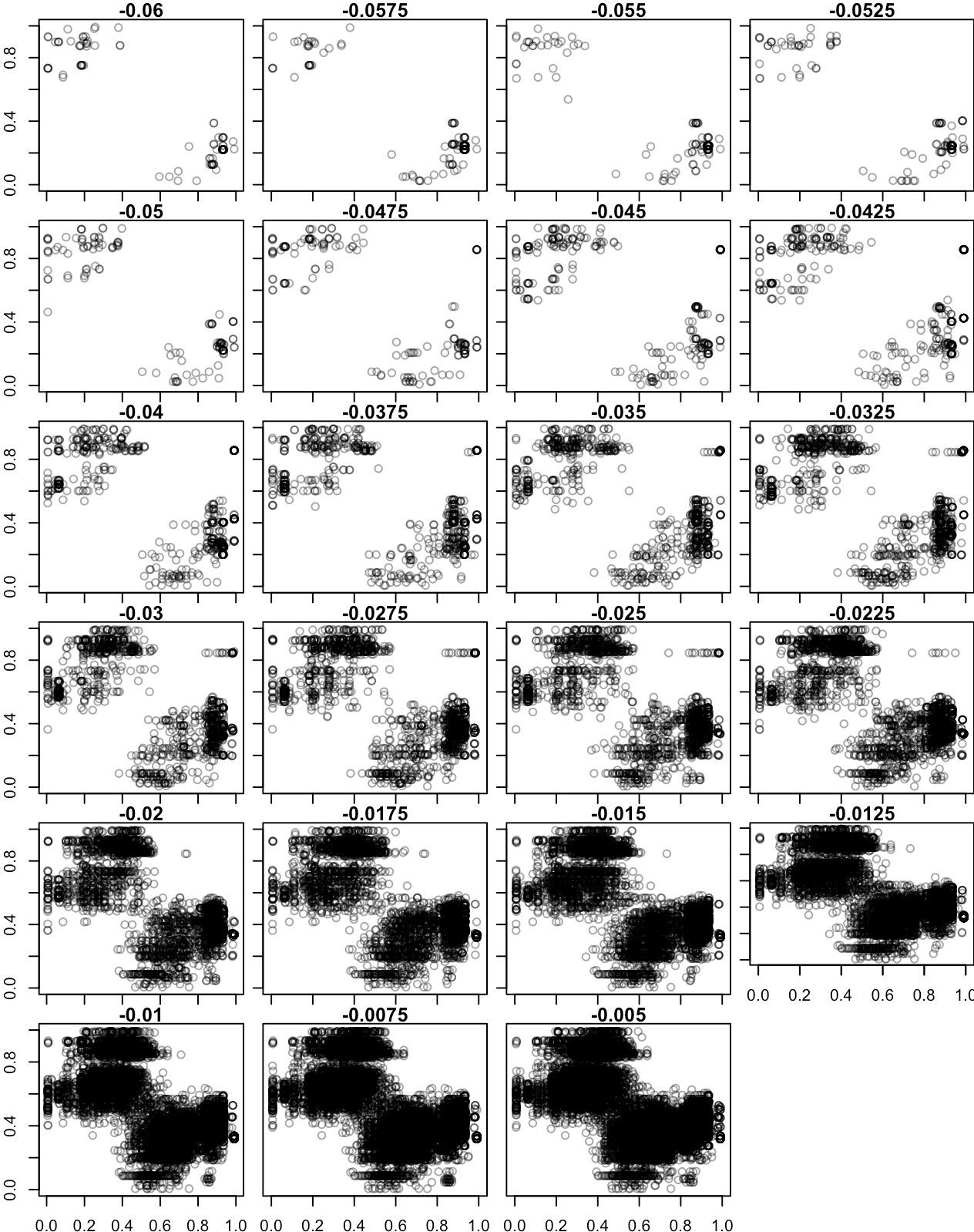

**Figure S13.** Hybrid index for candidate BDMI region pairs. Plot titles are X(2) cutoff values. Stricter (more negative) cutoffs require stronger ancestry distortion between candidate BDMI windows. X axis is the HI of region one of a BDMI pair and the y axis is the HI of region 2 of the BDMI pair.

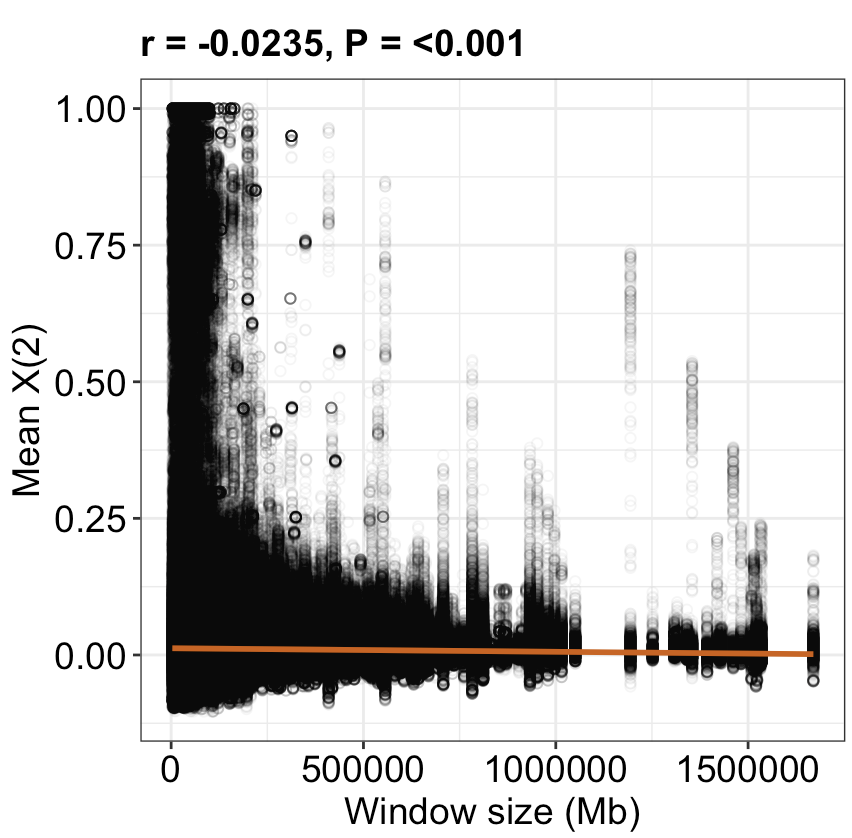

**Figure S14.** Window size effect on X(2) values in the AD scan analysis. Window sizes vary due to using an equal number of AIMs per window. There is a weak but significant correlation between window size and mean X(2) values.

**Supporting Tables**

**Table S1.** Global estimates of the demographic history of *F. aquilonia* and *F. polyctena* inferred using gIMble under models of strict divergence (DIV), migration without divergence (MIG), and isolation with migration (IM). Gray shading indicates the best-fit model. Parameter estimates are scaled in absolute units, i.e. number of individuals (*N_e_*) and years (*T;* converted from generations to years assuming 2.5 generations per year). Migration rate (*m*) estimates correspond to M (=4*N_e_m_e_*) individuals per generation (forwards in time).

| **Model** | **Ancestral *Ne*** | ***F. aqui Ne*** | ***F. pol Ne*** | ***T* (years)** | ***m* (M)** | **ΔlnCL** |
| --- | --- | --- | --- | --- | --- | --- |
| DIV | 1.36E+05 | 7.57E+04 | 1.20E+05 | 2.19E+05 | - | -17872 |
| MIG *F. aqui --> F. pol* | - | 1.07E+05 | 5.81E+04 | - | 6.93E-06 (1.61) | -12682 |
| MIG *F. pol --> F. aqui* | - | 3.25E+04 | 1.27E+05 | - | 9.16E-06 (1.19) | -24912 |
| IM *F. aqui --> F. pol* | 1.16E+05 | 1.04E+05 | 7.24E+04 | 5.78E+05 | 5.19E-06 (1.50) | 0 |
| IM *F. pol --> F. aqui* | 1.12E+05 | 4.30E+04 | 1.45E+05 | 5.36E+05 | 6.28E-06 (1.08) | -1111 |

**Table S2.** Chi-Square test results assessing whether the number of barrier windows on each chromosome matches the genome-wide occurrence rate (3.1%) or shows enrichment. The Chi-Square test statistic (X²) and corresponding p-values are listed for each chromosome. No significant enrichment of barriers was observed on any chromosome.

| **Chromosome** | **Observed barrier fraction (%)** | **X^2^** | **df** | **p-value** |
| --- | --- | --- | --- | --- |
| 1 | 3.9 | 0.10 | 1 | 0.750 |
| 2 | 3.1 | 0.00 | 1 | 1.000 |
| 3 | 2.3 | 0.09 | 1 | 0.770 |
| 4 | 4.4 | 0.37 | 1 | 0.541 |
| 5 | 1.7 | 0.41 | 1 | 0.523 |
| 6 | 2.3 | 0.05 | 1 | 0.830 |
| 7 | 4.2 | 0.09 | 1 | 0.760 |
| 8 | 0.2 | 1.81 | 1 | 0.179 |
| 9 | 2.9 | 0.00 | 1 | 1.000 |
| 10 | 5.1 | 0.28 | 1 | 0.599 |
| 11 | 3.8 | 0.00 | 1 | 1.000 |
| 12 | 0.8 | 1.28 | 1 | 0.258 |
| 13 | 2.6 | 0.00 | 1 | 1.000 |
| 14 | 3.9 | 0.00 | 1 | 0.974 |
| 15 | 1.7 | 0.15 | 1 | 0.698 |
| 16 | 1.2 | 0.29 | 1 | 0.588 |
| 17 | 8.8 | 2.35 | 1 | 0.125 |
| 18 | 3.7 | 0.00 | 1 | 1.000 |
| 19 | 1.1 | 0.21 | 1 | 0.645 |
| 20 | 3.3 | 0.00 | 1 | 1.000 |
| 21 | 2.3 | 0.00 | 1 | 0.944 |
| 22 | 4.0 | 0.00 | 1 | 0.972 |
| 23 | 0.7 | 0.37 | 1 | 0.541 |
| 24 | 4.1 | 0.00 | 1 | 1.000 |
| 25 | 4.2 | 0.01 | 1 | 0.941 |
| 26 | 3.4 | 0.00 | 1 | 1.000 |
| 27 | 5.0 | 0.04 | 1 | 0.841 |

**Table S3.** Mean genetic diversity (π) and heterozygosity (*H*) within *F. aquilonia* and *F. polyctena*, and mean genetic divergence (d_xy_) and genetic differentiation (F_st_) between the species.

| ***F. aquilonia* π** | ***F. polyctena* π** | ***F. aquilonia* *H*** | ***F. polyctena* *H*** | **d_xy_** | **F_st_** |
| --- | --- | --- | --- | --- | --- |
| 0.00217 | 0.00272 | 0.00201 | 0.00251 | 0.00327 | 0.18356 |

**Table S4.** Metadata for samples used in SNP calling for the X(2) analysis. Sample and ID are unique sample identifiers. Sex is the morphological identity of the sample sex. Nest is the name of the nest the sample was collected from. Dev_stage is the stage of development at time of DNA extraction (pupa, young_adult/sexual, adult). Year is the year the sample was collected. Ploidy is the ploidy level of the sample (1 = haploid, 2 = diploid). Caste is the sample social caste (m = male drone, w = female worker, q = female queen). avg_DP is the mean sequencing depth for the sample (decimal marker is ,). Perc_dup is the sequencing duplication rate. Est library size

| sample | ID | sex | nest | dev_stage | year | ploidy | caste | avg_DP | map_rate |
| --- | --- | --- | --- | --- | --- | --- | --- | --- | --- |
| **a10** | **m012-2011** | **m** | **Katajikko** | **NA** | **2011** | **1** | **m** | **4,272** | **984** |
| **a100** | **m001-2005** | **m** | **FA6** | **adult** | **2005** | **1** | **m** | **3,117** | **988** |
| **a101** | **m025-2004** | **m** | **Nest2** | **adult** | **2004** | **1** | **m** | **5,321** | **989** |
| **a102** | **m002-2005** | **m** | **FA6** | **adult** | **2005** | **1** | **m** | **5,749** | **991** |
| **a103** | **m006-2014** | **m** | **FA15** | **NA** | **2014** | **1** | **m** | **4,248** | **989** |
| **a104** | **m026-2004** | **m** | **Nest2** | **adult** | **2004** | **1** | **m** | **5,250** | **989** |
| **a105** | **m001-2018** | **m** | **FA18** | **pupa** | **2018** | **1** | **m** | **4,744** | **987** |
| **a106** | **m027-2004** | **m** | **Nest2** | **adult** | **2004** | **1** | **m** | **3,875** | **990** |
| **a107** | **m007-2014** | **m** | **FA15** | **NA** | **2014** | **1** | **m** | **3,478** | **990** |
| **a108** | **m002-2018** | **m** | **FA24** | **adult** | **2018** | **1** | **m** | **5,515** | **988** |
| **a109** | **m028-2004** | **m** | **Nest2** | **adult** | **2004** | **1** | **m** | **4,684** | **989** |
| **a110** | **m003-2005** | **m** | **FA6** | **adult** | **2005** | **1** | **m** | **4,684** | **985** |
| **a111** | **m029-2004** | **m** | **Nest2** | **adult** | **2004** | **1** | **m** | **4,429** | **987** |
| **a112** | **m004-2005** | **m** | **FA10** | **adult** | **2005** | **1** | **m** | **5,017** | **856** |
| **a113** | **m030-2004** | **m** | **Nest4** | **adult** | **2004** | **1** | **m** | **4,703** | **987** |
| **a115** | **m005-2005** | **m** | **FA6** | **adult** | **2005** | **1** | **m** | **4,354** | **985** |
| **a119** | **m012-2014** | **m** | **FA17** | **NA** | **2014** | **1** | **m** | **3,029** | **844** |
| **a12** | **m017-2014** | **m** | **FA17** | **NA** | **2014** | **1** | **m** | **3,507** | **989** |
| **a120** | **m003-2018** | **m** | **FA15** | **pupa** | **2018** | **1** | **m** | **4,576** | **989** |
| **a121** | **m004-2018** | **m** | **FA24** | **adult** | **2018** | **1** | **m** | **5,163** | **984** |
| **a122** | **m031-2004** | **m** | **Nest4** | **adult** | **2004** | **1** | **m** | **5,133** | **985** |
| **a123** | **m014-2014** | **m** | **FA15** | **NA** | **2014** | **1** | **m** | **3,895** | **989** |
| **a124** | **m005-2018** | **m** | **FA18** | **pupa** | **2018** | **1** | **m** | **3,963** | **984** |
| **a125** | **m006-2005** | **m** | **FA6** | **adult** | **2005** | **1** | **m** | **5,958** | **993** |
| **a126** | **m032-2004** | **m** | **Nest2** | **adult** | **2004** | **1** | **m** | **4,633** | **979** |
| **a127** | **m007-2005** | **m** | **FA5** | **adult** | **2005** | **1** | **m** | **3,874** | **992** |
| **a128** | **m015-2014** | **m** | **FA17** | **NA** | **2014** | **1** | **m** | **5,849** | **985** |
| **a129** | **m016-2014** | **m** | **FA17** | **NA** | **2014** | **1** | **m** | **4,158** | **990** |
| **a13** | **m021-2014** | **m** | **FA17** | **NA** | **2014** | **1** | **m** | **3,504** | **988** |
| **a130** | **m008-2005** | **m** | **FA5** | **adult** | **2005** | **1** | **m** | **3,846** | **993** |
| **a131** | **m009-2005** | **m** | **FA6** | **adult** | **2005** | **1** | **m** | **5,906** | **993** |
| **a132** | **m018-2014** | **m** | **FA17** | **NA** | **2014** | **1** | **m** | **4,435** | **991** |
| **a133** | **m033-2004** | **m** | **Nest5** | **adult** | **2004** | **1** | **m** | **5,763** | **981** |
| **a135** | **m006-2018** | **m** | **FA18** | **pupa** | **2018** | **1** | **m** | **5,023** | **989** |
| **a136** | **m020-2014** | **m** | **FA15** | **NA** | **2014** | **1** | **m** | **4,364** | **990** |
| **a138** | **m010-2005** | **m** | **FA6** | **adult** | **2005** | **1** | **m** | **6,237** | **989** |
| **a139** | **m011-2005** | **m** | **FA6** | **adult** | **2005** | **1** | **m** | **3,401** | **992** |
| **a141** | **m035-2004** | **m** | **Nest5** | **adult** | **2004** | **1** | **m** | **4,303** | **988** |
| **a142** | **m012-2005** | **m** | **FA6** | **adult** | **2005** | **1** | **m** | **5,557** | **992** |
| **a145** | **m023-2014** | **m** | **FA17** | **NA** | **2014** | **1** | **m** | **5,176** | **982** |
| **a147** | **m013-2005** | **m** | **FA6** | **adult** | **2005** | **1** | **m** | **3,122** | **991** |
| **a148** | **m009-2018** | **m** | **FA39** | **pupa** | **2018** | **1** | **m** | **4,412** | **990** |
| **a149** | **m037-2004** | **m** | **Nest3** | **adult** | **2004** | **1** | **m** | **5,029** | **989** |
| **a15** | **m016-2018** | **m** | **FA15** | **pupa** | **2018** | **1** | **m** | **4,039** | **983** |
| **a150** | **m038-2004** | **m** | **Nest5** | **adult** | **2004** | **1** | **m** | **5,575** | **984** |
| **a151** | **m039-2004** | **m** | **Nest5** | **adult** | **2004** | **1** | **m** | **5,173** | **989** |
| **a152** | **m024-2014** | **m** | **FA15** | **NA** | **2014** | **1** | **m** | **4,226** | **988** |
| **a153** | **m025-2014** | **m** | **FA15** | **NA** | **2014** | **1** | **m** | **5,279** | **986** |
| **a155** | **m012-2018** | **m** | **FA38** | **pupa** | **2018** | **1** | **m** | **5,879** | **983** |
| **a157** | **m014-2018** | **m** | **FA39** | **pupa** | **2018** | **1** | **m** | **4,420** | **991** |
| **a158** | **m040-2004** | **m** | **Nest4** | **adult** | **2004** | **1** | **m** | **5,243** | **980** |
| **a159** | **m015-2018** | **m** | **FA38** | **pupa** | **2018** | **1** | **m** | **4,658** | **985** |
| **a16** | **m019-2018** | **m** | **FA24** | **adult** | **2018** | **1** | **m** | **4,432** | **987** |
| **a160** | **m014-2005** | **m** | **FA10** | **adult** | **2005** | **1** | **m** | **4,335** | **989** |
| **a161** | **m015-2005** | **m** | **FA6** | **adult** | **2005** | **1** | **m** | **3,694** | **983** |
| **a162** | **m016-2005** | **m** | **FA5** | **adult** | **2005** | **1** | **m** | **3,258** | **987** |
| **a163** | **m026-2014** | **m** | **FA17** | **NA** | **2014** | **1** | **m** | **4,352** | **989** |
| **a164** | **m017-2018** | **m** | **FA38** | **pupa** | **2018** | **1** | **m** | **5,244** | **987** |
| **a165** | **m041-2004** | **m** | **Nest4** | **adult** | **2004** | **1** | **m** | **5,179** | **989** |
| **a167** | **m018-2018** | **m** | **FA24** | **adult** | **2018** | **1** | **m** | **4,941** | **977** |
| **a168** | **m042-2004** | **m** | **Nest2** | **adult** | **2004** | **1** | **m** | **4,533** | **992** |
| **a169** | **m043-2004** | **m** | **Nest2** | **adult** | **2004** | **1** | **m** | **6,200** | **986** |
| **a17** | **m032-2014** | **m** | **FA15** | **NA** | **2014** | **1** | **m** | **3,430** | **989** |
| **a172** | **m021-2018** | **m** | **FA39** | **pupa** | **2018** | **1** | **m** | **6,113** | **987** |
| **a173** | **m017-2005** | **m** | **FA6** | **adult** | **2005** | **1** | **m** | **3,578** | **992** |
| **a175** | **m022-2018** | **m** | **FA39** | **pupa** | **2018** | **1** | **m** | **3,924** | **987** |
| **a176** | **m030-2014** | **m** | **FA17** | **NA** | **2014** | **1** | **m** | **4,718** | **978** |
| **a177** | **m044-2004** | **m** | **Nest4** | **adult** | **2004** | **1** | **m** | **4,800** | **970** |
| **a179** | **m031-2014** | **m** | **FA15** | **NA** | **2014** | **1** | **m** | **4,753** | **986** |
| **a18** | **m048-2004** | **m** | **Nest4** | **adult** | **2004** | **1** | **m** | **4,655** | **987** |
| **a180** | **m018-2005** | **m** | **FA5** | **adult** | **2005** | **1** | **m** | **4,718** | **989** |
| **a181** | **m046-2004** | **m** | **Nest3** | **adult** | **2004** | **1** | **m** | **4,756** | **985** |
| **a182** | **m033-2014** | **m** | **FA17** | **NA** | **2014** | **1** | **m** | **3,480** | **987** |
| **a184** | **m034-2014** | **m** | **FA17** | **NA** | **2014** | **1** | **m** | **4,698** | **934** |
| **a186** | **m019-2005** | **m** | **FA5** | **adult** | **2005** | **1** | **m** | **3,849** | **989** |
| **a187** | **m020-2005** | **m** | **FA5** | **adult** | **2005** | **1** | **m** | **4,547** | **987** |
| **a188** | **m024-2018** | **m** | **FA24** | **adult** | **2018** | **1** | **m** | **5,282** | **987** |
| **a189** | **m021-2005** | **m** | **FA6** | **adult** | **2005** | **1** | **m** | **5,571** | **991** |
| **a19** | **m035-2014** | **m** | **FA17** | **NA** | **2014** | **1** | **m** | **3,451** | **964** |
| **a190** | **m049-2004** | **m** | **Nest4** | **adult** | **2004** | **1** | **m** | **5,830** | **986** |
| **a191** | **m022-2005** | **m** | **FA6** | **adult** | **2005** | **1** | **m** | **4,788** | **991** |
| **a192** | **m023-2005** | **m** | **FA6** | **adult** | **2005** | **1** | **m** | **3,482** | **991** |
| **a20** | **m025-2018** | **m** | **FA18** | **pupa** | **2018** | **1** | **m** | **4,647** | **993** |
| **a21** | **m051-2004** | **m** | **Nest2** | **adult** | **2004** | **1** | **m** | **4,249** | **988** |
| **a22** | **m026-2018** | **m** | **FA39** | **pupa** | **2018** | **1** | **m** | **4,250** | **987** |
| **a24** | **m024-2005** | **m** | **FA5** | **adult** | **2005** | **1** | **m** | **3,907** | **991** |
| **a25** | **m036-2014** | **m** | **FA17** | **NA** | **2014** | **1** | **m** | **4,532** | **988** |
| **a28** | **m028-2018** | **m** | **FA24** | **adult** | **2018** | **1** | **m** | **3,531** | **977** |
| **a29** | **m025-2005** | **m** | **FA10** | **adult** | **2005** | **1** | **m** | **3,776** | **989** |
| **a3** | **m059-2004** | **m** | **Nest3** | **adult** | **2004** | **1** | **m** | **3,552** | **990** |
| **a30** | **m029-2018** | **m** | **FA38** | **pupa** | **2018** | **1** | **m** | **3,986** | **987** |
| **a32** | **m030-2018** | **m** | **FA38** | **pupa** | **2018** | **1** | **m** | **4,800** | **986** |
| **a33** | **m037-2014** | **m** | **FA15** | **NA** | **2014** | **1** | **m** | **3,729** | **988** |
| **a35** | **m026-2005** | **m** | **FA10** | **adult** | **2005** | **1** | **m** | **4,088** | **985** |
| **a36** | **m027-2005** | **m** | **FA6** | **adult** | **2005** | **1** | **m** | **3,466** | **991** |
| **a37** | **m057-2004** | **m** | **Nest4** | **adult** | **2004** | **1** | **m** | **3,985** | **984** |
| **a39** | **m028-2005** | **m** | **FA10** | **adult** | **2005** | **1** | **m** | **3,392** | **989** |
| **a4** | **m065-2004** | **m** | **Nest5** | **adult** | **2004** | **1** | **m** | **6,171** | **985** |
| **a40** | **m060-2004** | **m** | **Nest5** | **adult** | **2004** | **1** | **m** | **6,223** | **984** |
| **a41** | **m038-2014** | **m** | **FA15** | **NA** | **2014** | **1** | **m** | **4,363** | **990** |
| **a44** | **m031-2018** | **m** | **FA18** | **pupa** | **2018** | **1** | **m** | **4,393** | **991** |
| **a45** | **m063-2004** | **m** | **Nest2** | **adult** | **2004** | **1** | **m** | **4,315** | **990** |
| **a46** | **m039-2014** | **m** | **FA17** | **NA** | **2014** | **1** | **m** | **3,899** | **844** |
| **a47** | **m040-2014** | **m** | **FA17** | **NA** | **2014** | **1** | **m** | **5,190** | **985** |
| **a48** | **m064-2004** | **m** | **Nest3** | **adult** | **2004** | **1** | **m** | **3,866** | **986** |
| **a49** | **m029-2005** | **m** | **FA6** | **adult** | **2005** | **1** | **m** | **4,328** | **988** |
| **a5** | **m032-2005** | **m** | **FA5** | **adult** | **2005** | **1** | **m** | **4,010** | **990** |
| **a51** | **m030-2005** | **m** | **FA10** | **adult** | **2005** | **1** | **m** | **3,701** | **988** |
| **a52** | **m066-2004** | **m** | **Nest4** | **adult** | **2004** | **1** | **m** | **4,737** | **984** |
| **a53** | **m031-2005** | **m** | **FA6** | **adult** | **2005** | **1** | **m** | **3,394** | **985** |
| **a55** | **m032-2018** | **m** | **FA18** | **pupa** | **2018** | **1** | **m** | **4,929** | **985** |
| **a56** | **m033-2018** | **m** | **FA38** | **pupa** | **2018** | **1** | **m** | **5,163** | **986** |
| **a57** | **m068-2004** | **m** | **Nest2** | **adult** | **2004** | **1** | **m** | **5,023** | **988** |
| **a58** | **m034-2018** | **m** | **FA15** | **pupa** | **2018** | **1** | **m** | **5,189** | **989** |
| **a59** | **m042-2014** | **m** | **FA17** | **NA** | **2014** | **1** | **m** | **3,560** | **983** |
| **a6** | **m035-2005** | **m** | **FA10** | **adult** | **2005** | **1** | **m** | **3,552** | **991** |
| **a60** | **m033-2005** | **m** | **FA10** | **adult** | **2005** | **1** | **m** | **4,691** | **987** |
| **a61** | **m035-2018** | **m** | **FA25** | **pupa** | **2018** | **1** | **m** | **4,296** | **988** |
| **a62** | **m069-2004** | **m** | **Nest3** | **adult** | **2004** | **1** | **m** | **4,979** | **986** |
| **a63** | **m070-2004** | **m** | **Nest2** | **adult** | **2004** | **1** | **m** | **3,974** | **989** |
| **a64** | **m043-2014** | **m** | **FA17** | **NA** | **2014** | **1** | **m** | **4,614** | **901** |
| **a66** | **m036-2018** | **m** | **FA24** | **adult** | **2018** | **1** | **m** | **4,670** | **980** |
| **a67** | **m045-2014** | **m** | **FA15** | **NA** | **2014** | **1** | **m** | **3,438** | **991** |
| **a68** | **m034-2005** | **m** | **FA6** | **adult** | **2005** | **1** | **m** | **4,119** | **989** |
| **a69** | **m046-2014** | **m** | **FA15** | **NA** | **2014** | **1** | **m** | **4,003** | **990** |
| **a70** | **m047-2014** | **m** | **FA17** | **NA** | **2014** | **1** | **m** | **3,940** | **987** |
| **a71** | **m037-2018** | **m** | **FA24** | **adult** | **2018** | **1** | **m** | **3,958** | **979** |
| **a72** | **m071-2004** | **m** | **Nest5** | **adult** | **2004** | **1** | **m** | **4,194** | **983** |
| **a73** | **m038-2018** | **m** | **FA15** | **pupa** | **2018** | **1** | **m** | **4,292** | **989** |
| **a74** | **m072-2004** | **m** | **Nest4** | **adult** | **2004** | **1** | **m** | **6,465** | **986** |
| **a75** | **m073-2004** | **m** | **Nest2** | **adult** | **2004** | **1** | **m** | **5,091** | **990** |
| **a76** | **m036-2005** | **m** | **FA5** | **adult** | **2005** | **1** | **m** | **5,296** | **987** |
| **a81** | **m039-2018** | **m** | **FA25** | **pupa** | **2018** | **1** | **m** | **4,050** | **991** |
| **a82** | **m040-2018** | **m** | **FA25** | **pupa** | **2018** | **1** | **m** | **4,672** | **989** |
| **a83** | **m076-2004** | **m** | **Nest4** | **adult** | **2004** | **1** | **m** | **5,180** | **983** |
| **a84** | **m077-2004** | **m** | **Nest4** | **adult** | **2004** | **1** | **m** | **4,734** | **986** |
| **a85** | **m078-2004** | **m** | **Nest5** | **adult** | **2004** | **1** | **m** | **4,671** | **979** |
| **a86** | **m050-2014** | **m** | **FA17** | **NA** | **2014** | **1** | **m** | **4,756** | **811** |
| **a87** | **m037-2005** | **m** | **FA6** | **adult** | **2005** | **1** | **m** | **4,009** | **993** |
| **a88** | **m079-2004** | **m** | **Nest4** | **adult** | **2004** | **1** | **m** | **4,838** | **988** |
| **a89** | **m080-2004** | **m** | **Nest4** | **adult** | **2004** | **1** | **m** | **4,628** | **988** |
| **a9** | **m014-2011** | **m** | **FA12** | **NA** | **2011** | **1** | **m** | **4,293** | **984** |
| **a90** | **m041-2018** | **m** | **FA24** | **adult** | **2018** | **1** | **m** | **5,961** | **987** |
| **a91** | **m038-2005** | **m** | **FA5** | **adult** | **2005** | **1** | **m** | **3,214** | **986** |
| **a92** | **m051-2014** | **m** | **FA17** | **NA** | **2014** | **1** | **m** | **4,034** | **977** |
| **a94** | **m053-2014** | **m** | **FA15** | **NA** | **2014** | **1** | **m** | **4,310** | **990** |
| **a96** | **m039-2005** | **m** | **FA5** | **adult** | **2005** | **1** | **m** | **4,414** | **987** |
| **a98** | **m040-2005** | **m** | **FA5** | **adult** | **2005** | **1** | **m** | **4,580** | **989** |
| **a99** | **m081-2004** | **m** | **Nest2** | **adult** | **2004** | **1** | **m** | **5,703** | **990** |
| **FA04_3m** | **m045-2018** | **m** | **FA04** | **adult** | **2018** | **1** | **m** | **4,064** | **977** |
| **FA07_1m** | **m051-2018** | **m** | **FA07** | **adult** | **2018** | **1** | **m** | **4,009** | **984** |
| **FA12_1m** | **m060-2018** | **m** | **FA12** | **adult** | **2018** | **1** | **m** | **4,735** | **981** |
| **FA12_2m** | **m061-2018** | **m** | **FA12** | **adult** | **2018** | **1** | **m** | **4,316** | **973** |
| **FA12_3m** | **m062-2018** | **m** | **FA12** | **adult** | **2018** | **1** | **m** | **4,340** | **981** |
| **FA12_4m** | **m063-2018** | **m** | **FA12** | **adult** | **2018** | **1** | **m** | **5,307** | **979** |
| **FA12_5m** | **m064-2018** | **m** | **FA12** | **adult** | **2018** | **1** | **m** | **4,492** | **981** |
| **FA12_6m** | **m065-2018** | **m** | **FA12** | **adult** | **2018** | **1** | **m** | **5,372** | **984** |
| **FA12_7m** | **m066-2018** | **m** | **FA12** | **adult** | **2018** | **1** | **m** | **5,225** | **983** |
| **FA15_1m** | **m069-2018** | **m** | **FA15** | **adult** | **2018** | **1** | **m** | **5,328** | **985** |
| **FA15_3m** | **m070-2018** | **m** | **FA15** | **adult** | **2018** | **1** | **m** | **5,273** | **980** |
| **FA16_1m** | **m073-2018** | **m** | **FA16** | **adult** | **2018** | **1** | **m** | **4,262** | **985** |
| **FA16_2m** | **m074-2018** | **m** | **FA16** | **adult** | **2018** | **1** | **m** | **4,231** | **983** |
| **FA16_3m** | **m075-2018** | **m** | **FA16** | **adult** | **2018** | **1** | **m** | **4,535** | **984** |
| **FAu14_1m** | **m079-2018** | **m** | **FAu14** | **adult** | **2018** | **1** | **m** | **3,631** | **983** |
| **s193** | **m020-2021** | **m** | **FA15** | **adult_sexual** | **2021** | **1** | **m** | **3,787** | **982** |
| **s194** | **m021-2021** | **m** | **FA15** | **adult_sexual** | **2021** | **1** | **m** | **3,965** | **980** |
| **s196** | **m022-2020** | **m** | **FA2014_2** | **young_adult** | **2020** | **1** | **m** | **4,156** | **984** |
| **s197** | **m023-2020** | **m** | **FA39** | **young_adult** | **2020** | **1** | **m** | **4,601** | **982** |
| **s198** | **m024-2020** | **m** | **FA2014_2** | **young_adult** | **2020** | **1** | **m** | **4,755** | **974** |
| **s201** | **m025-2020** | **m** | **FA39** | **young_adult** | **2020** | **1** | **m** | **5,082** | **982** |
| **s205** | **m001-2011** | **m** | **Katajikko** | **adult** | **2011** | **1** | **m** | **3,587** | **949** |
| **s209** | **m027-2020** | **m** | **FA2014_2** | **young_adult** | **2020** | **1** | **m** | **4,839** | **984** |
| **s212** | **m002-2011** | **m** | **Katajikko** | **adult** | **2011** | **1** | **m** | **3,907** | **937** |
| **s213** | **m028-2020** | **m** | **FA2020** | **young_adult** | **2020** | **1** | **m** | **4,233** | **984** |
| **s214** | **m022-2021** | **m** | **FA15** | **adult_sexual** | **2021** | **1** | **m** | **3,800** | **981** |
| **s215** | **m023-2021** | **m** | **FA38** | **adult_sexual** | **2021** | **1** | **m** | **4,375** | **988** |
| **s216** | **m024-2021** | **m** | **FA20** | **adult_sexual** | **2021** | **1** | **m** | **4,099** | **977** |
| **s217** | **m025-2021** | **m** | **FAuus2014A2** | **adult_sexual** | **2021** | **1** | **m** | **2,773** | **994** |
| **s222** | **m026-2021** | **m** | **FA38** | **adult_sexual** | **2021** | **1** | **m** | **3,943** | **982** |
| **s223** | **m029-2020** | **m** | **FA39** | **young_adult** | **2020** | **1** | **m** | **4,332** | **986** |
| **s224** | **m030-2020** | **m** | **FA2020** | **young_adult** | **2020** | **1** | **m** | **4,451** | **984** |
| **s226** | **m027-2021** | **m** | **FA15** | **adult_sexual** | **2021** | **1** | **m** | **4,413** | **979** |
| **s227** | **m028-2021** | **m** | **FA13** | **adult_sexual** | **2021** | **1** | **m** | **4,157** | **974** |
| **s228** | **m031-2020** | **m** | **FA2014_2** | **young_adult** | **2020** | **1** | **m** | **4,210** | **979** |
| **s232** | **m029-2021** | **m** | **FA20** | **adult_sexual** | **2021** | **1** | **m** | **3,443** | **986** |
| **s233** | **m030-2021** | **m** | **FA15** | **adult_sexual** | **2021** | **1** | **m** | **4,303** | **986** |
| **s234** | **m031-2021** | **m** | **FA15** | **adult_sexual** | **2021** | **1** | **m** | **4,237** | **980** |
| **s235** | **m003-2011** | **m** | **Katajikko** | **adult** | **2011** | **1** | **m** | **3,904** | **982** |
| **s240** | **m033-2021** | **m** | **FA38** | **adult_sexual** | **2021** | **1** | **m** | **4,326** | **986** |
| **s244** | **m033-2020** | **m** | **FA33** | **young_adult** | **2020** | **1** | **m** | **4,084** | **980** |
| **s248** | **m005-2011** | **m** | **Katajikko** | **adult** | **2011** | **1** | **m** | **3,221** | **986** |
| **s249** | **m034-2020** | **m** | **FA2014_2** | **young_adult** | **2020** | **1** | **m** | **4,624** | **981** |
| **s250** | **m035-2020** | **m** | **FA2020** | **young_adult** | **2020** | **1** | **m** | **4,261** | **981** |
| **s251** | **m036-2020** | **m** | **FA39** | **young_adult** | **2020** | **1** | **m** | **4,770** | **981** |
| **s252** | **m035-2021** | **m** | **FA38** | **adult_sexual** | **2021** | **1** | **m** | **3,077** | **972** |
| **s253** | **m036-2021** | **m** | **FA38** | **adult_sexual** | **2021** | **1** | **m** | **3,783** | **983** |
| **s255** | **m037-2020** | **m** | **FA2020** | **young_adult** | **2020** | **1** | **m** | **4,323** | **984** |
| **s257** | **m039-2020** | **m** | **FA2020** | **young_adult** | **2020** | **1** | **m** | **4,630** | **983** |
| **s259** | **m040-2020** | **m** | **FA33** | **young_adult** | **2020** | **1** | **m** | **4,108** | **984** |
| **s261** | **m006-2011** | **m** | **Pohjrannw** | **NA** | **2011** | **1** | **m** | **2,416** | **985** |
| **s266** | **m038-2021** | **m** | **FA20** | **adult_sexual** | **2021** | **1** | **m** | **4,690** | **976** |
| **s267** | **m007-2011** | **m** | **FA12** | **adult** | **2011** | **1** | **m** | **4,702** | **984** |
| **s272** | **m041-2020** | **m** | **FA2020** | **young_adult** | **2020** | **1** | **m** | **5,002** | **984** |
| **s273** | **m040-2021** | **m** | **FAuus2014A2** | **adult_sexual** | **2021** | **1** | **m** | **5,027** | **981** |
| **s275** | **m041-2021** | **m** | **FA38** | **adult_sexual** | **2021** | **1** | **m** | **4,698** | **985** |
| **s277** | **m042-2021** | **m** | **FAuus2014A2** | **adult_sexual** | **2021** | **1** | **m** | **3,591** | **952** |
| **s278** | **m043-2021** | **m** | **FA38** | **adult_sexual** | **2021** | **1** | **m** | **3,883** | **946** |
| **s279** | **m044-2021** | **m** | **FA38** | **adult_sexual** | **2021** | **1** | **m** | **4,186** | **980** |
| **s281** | **m042-2020** | **m** | **FA33** | **young_adult** | **2020** | **1** | **m** | **4,669** | **981** |
| **s284** | **m008-2011** | **m** | **FA12** | **adult** | **2011** | **1** | **m** | **3,636** | **983** |
| **s285** | **m043-2020** | **m** | **FA2020** | **young_adult** | **2020** | **1** | **m** | **4,269** | **980** |
| **s288** | **m045-2021** | **m** | **FA15** | **adult_sexual** | **2021** | **1** | **m** | **3,327** | **781** |
| **s289** | **m044-2020** | **m** | **FA2014_2** | **young_adult** | **2020** | **1** | **m** | **4,804** | **979** |
| **s291** | **m013-2021** | **m** | **FAuus2014A2** | **adult_sexual** | **2021** | **1** | **m** | **9,576** | **953** |
| **s294** | **m046-2021** | **m** | **FA15** | **adult_sexual** | **2021** | **1** | **m** | **3,676** | **979** |
| **s297** | **m045-2020** | **m** | **FA39** | **young_adult** | **2020** | **1** | **m** | **5,281** | **975** |
| **s300** | **m046-2020** | **m** | **FA39** | **young_adult** | **2020** | **1** | **m** | **5,264** | **975** |
| **s301** | **m047-2021** | **m** | **FAuus2014A2** | **adult_sexual** | **2021** | **1** | **m** | **4,774** | **988** |
| **s302** | **m047-2020** | **m** | **FA39** | **young_adult** | **2020** | **1** | **m** | **3,786** | **613** |
| **s304** | **m048-2020** | **m** | **FA2014_2** | **young_adult** | **2020** | **1** | **m** | **4,662** | **981** |
| **s305** | **m009-2011** | **m** | **Katajikko** | **adult** | **2011** | **1** | **m** | **3,847** | **985** |
| **s307** | **m049-2020** | **m** | **FA2014_2** | **young_adult** | **2020** | **1** | **m** | **3,951** | **811** |
| **s309** | **m050-2020** | **m** | **FA2020** | **young_adult** | **2020** | **1** | **m** | **4,414** | **983** |
| **s312** | **m051-2020** | **m** | **FA39** | **young_adult** | **2020** | **1** | **m** | **4,562** | **977** |
| **s315** | **m010-2011** | **m** | **Katajikko** | **adult** | **2011** | **1** | **m** | **4,397** | **983** |
| **s316** | **m052-2020** | **m** | **FA39** | **young_adult** | **2020** | **1** | **m** | **5,070** | **979** |
| **s318** | **m053-2020** | **m** | **FA39** | **young_adult** | **2020** | **1** | **m** | **4,753** | **980** |
| **s319** | **m048-2021** | **m** | **FAuus2014A2** | **adult_sexual** | **2021** | **1** | **m** | **4,582** | **977** |
| **s321** | **m049-2021** | **m** | **FA20** | **adult_sexual** | **2021** | **1** | **m** | **3,970** | **974** |
| **s322** | **m050-2021** | **m** | **FA38** | **adult_sexual** | **2021** | **1** | **m** | **4,059** | **981** |
| **s325** | **m051-2021** | **m** | **FA20** | **adult_sexual** | **2021** | **1** | **m** | **4,234** | **985** |
| **s327** | **m052-2021** | **m** | **FA15** | **adult_sexual** | **2021** | **1** | **m** | **4,247** | **981** |
| **s330** | **m053-2021** | **m** | **FA20** | **adult_sexual** | **2021** | **1** | **m** | **4,717** | **987** |
| **s335** | **m054-2020** | **m** | **FA39** | **young_adult** | **2020** | **1** | **m** | **4,640** | **973** |
| **s337** | **m055-2021** | **m** | **FA38** | **adult_sexual** | **2021** | **1** | **m** | **4,317** | **985** |
| **s339** | **m056-2021** | **m** | **FA38** | **adult_sexual** | **2021** | **1** | **m** | **4,142** | **987** |
| **s341** | **m055-2020** | **m** | **FA2020** | **young_adult** | **2020** | **1** | **m** | **4,635** | **985** |
| **s342** | **m057-2021** | **m** | **FA20** | **adult_sexual** | **2021** | **1** | **m** | **3,089** | **983** |
| **s344** | **m058-2021** | **m** | **FA20** | **adult_sexual** | **2021** | **1** | **m** | **3,310** | **985** |
| **s346** | **m056-2020** | **m** | **FA2014_2** | **young_adult** | **2020** | **1** | **m** | **4,489** | **982** |
| **s347** | **m057-2020** | **m** | **FA2020** | **young_adult** | **2020** | **1** | **m** | **4,573** | **983** |
| **s348** | **m059-2021** | **m** | **FAuus2014A2** | **adult_sexual** | **2021** | **1** | **m** | **2,970** | **953** |
| **s349** | **m058-2020** | **m** | **FA2020** | **young_adult** | **2020** | **1** | **m** | **4,773** | **985** |
| **s350** | **m059-2020** | **m** | **FA2014_2** | **young_adult** | **2020** | **1** | **m** | **4,891** | **982** |
| **s351** | **m060-2020** | **m** | **FA2020** | **young_adult** | **2020** | **1** | **m** | **3,939** | **970** |
| **s352** | **m061-2020** | **m** | **FA2014_2** | **young_adult** | **2020** | **1** | **m** | **4,445** | **978** |
| **RN356** | **FA33_09m** | **m** | **FA33** | **adult** | **2022** | **1** | **m** | **4,880** | **973** |
| **RN359** | **Bermuda21_10m** | **m** | **Bermuda21** | **adult** | **2022** | **1** | **m** | **4,597** | **975** |
| **RN360** | **FA33_04m** | **m** | **FA33** | **adult** | **2022** | **1** | **m** | **4,437** | **980** |
| **RN362** | **FA23_10m** | **m** | **FA23** | **adult** | **2022** | **1** | **m** | **4,618** | **976** |
| **RN363** | **FA12_05m** | **m** | **FA12** | **adult** | **2022** | **1** | **m** | **4,054** | **978** |
| **RN364** | **FA33_07m** | **m** | **FA33** | **adult** | **2022** | **1** | **m** | **3,340** | **981** |
| **RN365** | **FA23_04m** | **m** | **FA23** | **adult** | **2022** | **1** | **m** | **2,736** | **964** |
| **RN367** | **Bermuda21_01m** | **m** | **Bermuda21** | **adult** | **2022** | **1** | **m** | **2,897** | **974** |
| **RN368** | **FA12_04m** | **m** | **FA12** | **adult** | **2022** | **1** | **m** | **3,078** | **982** |
| **RN369** | **FA23_01m** | **m** | **FA23** | **adult** | **2022** | **1** | **m** | **2,799** | **973** |
| **RN370** | **FA12_03m** | **m** | **FA12** | **adult** | **2022** | **1** | **m** | **2,703** | **983** |
| **RN372** | **FA12_10m** | **m** | **FA12** | **adult** | **2022** | **1** | **m** | **2,434** | **980** |
| **RN375** | **FA23_03m** | **m** | **FA23** | **adult** | **2022** | **1** | **m** | **3,803** | **970** |
| **RN376** | **FA12_01m** | **m** | **FA12** | **adult** | **2022** | **1** | **m** | **3,797** | **981** |
| **RN378** | **FA33_03m** | **m** | **FA33** | **adult** | **2022** | **1** | **m** | **2,383** | **982** |
| **RN379** | **Bermuda21_06m** | **m** | **Bermuda21** | **adult** | **2022** | **1** | **m** | **5,227** | **966** |
| **RN381** | **Bermuda21_04m** | **m** | **Bermuda21** | **adult** | **2022** | **1** | **m** | **4,373** | **975** |
| **RN382** | **FA33_01m** | **m** | **FA33** | **adult** | **2022** | **1** | **m** | **4,413** | **978** |
| **RN384** | **FA33_08m** | **m** | **FA33** | **adult** | **2022** | **1** | **m** | **4,524** | **976** |
| **RN385** | **Bermuda21_05m** | **m** | **Bermuda21** | **adult** | **2022** | **1** | **m** | **4,197** | **978** |
| **RN390** | **FA23_09m** | **m** | **FA23** | **adult** | **2022** | **1** | **m** | **4,040** | **976** |
| **RN391** | **FA12_08m** | **m** | **FA12** | **adult** | **2022** | **1** | **m** | **4,361** | **976** |
| **RN393** | **FA12_07m** | **m** | **FA12** | **adult** | **2022** | **1** | **m** | **3,978** | **980** |
| **RN396** | **Bermuda21_08m** | **m** | **Bermuda21** | **adult** | **2022** | **1** | **m** | **3,709** | **974** |
| **RN397** | **Bermuda21_09m** | **m** | **Bermuda21** | **adult** | **2022** | **1** | **m** | **3,412** | **979** |
| **RN398** | **FA12_02m** | **m** | **FA12** | **adult** | **2022** | **1** | **m** | **3,429** | **977** |
| **RN399** | **FA33_10m** | **m** | **FA33** | **adult** | **2022** | **1** | **m** | **3,751** | **977** |
| **RN400** | **FA23_08m** | **m** | **FA23** | **adult** | **2022** | **1** | **m** | **3,836** | **977** |
| **RN401** | **FA33_02m** | **m** | **FA33** | **adult** | **2022** | **1** | **m** | **3,455** | **978** |
| **RN403** | **FA23_06m** | **m** | **FA23** | **adult** | **2022** | **1** | **m** | **3,672** | **968** |
| **RN404** | **FA23_07m** | **m** | **FA23** | **adult** | **2022** | **1** | **m** | **2,455** | **966** |
| **RN405** | **FA12_06m** | **m** | **FA12** | **adult** | **2022** | **1** | **m** | **2,761** | **964** |
| **RN406** | **FA23_02m** | **m** | **FA23** | **adult** | **2022** | **1** | **m** | **2,263** | **967** |
| **RN407** | **FA12_09m** | **m** | **FA12** | **adult** | **2022** | **1** | **m** | **2,729** | **979** |
| **RN408** | **FA23_05m** | **m** | **FA23** | **adult** | **2022** | **1** | **m** | **2,716** | **971** |
| **RN409** | **FA33_05m** | **m** | **FA33** | **adult** | **2022** | **1** | **m** | **2,478** | **981** |
| **RN411** | **Bermuda21_07m** | **m** | **Bermuda21** | **adult** | **2022** | **1** | **m** | **3,990** | **972** |
| **RN412** | **FA33_06m** | **m** | **FA33** | **adult** | **2022** | **1** | **m** | **3,472** | **976** |
| **RN413** | **Bermuda21_02m** | **m** | **Bermuda21** | **adult** | **2022** | **1** | **m** | **3,363** | **975** |
| **RN414** | **Bermuda21_03m** | **m** | **Bermuda21** | **adult** | **2022** | **1** | **m** | **2,896** | **974** |

**Table S5**. BDMI count and coverage data for the X(2) analysis.

| **cutoff** | **total_dmis** | **intra_pairs** | **intra_perc** | **inter_pairs** | **inter_perc** | **total_bp** | **total_frac_genome** | **mean_region_size** | **min_region_size** | **max_region_size** |
| --- | --- | --- | --- | --- | --- | --- | --- | --- | --- | --- |
| -0.06 | 87 | 3 | 0.034 | 84 | 0.966 | 4250006 | 0.02 | 77972 | 3722 | 418576 |
| -0.0575 | 118 | 3 | 0.025 | 115 | 0.975 | 4974634 | 0.023 | 66804 | 3722 | 782744 |
| -0.055 | 122 | 2 | 0.016 | 120 | 0.984 | 5046008 | 0.024 | 63783 | 3722 | 707440 |
| -0.0525 | 118 | 4 | 0.034 | 114 | 0.966 | 7989334 | 0.038 | 100861 | 4485 | 1520567 |
| -0.05 | 121 | 4 | 0.033 | 117 | 0.967 | 8190308 | 0.038 | 104576 | 3722 | 932927 |
| -0.0475 | 158 | 7 | 0.044 | 151 | 0.956 | 12905987 | 0.061 | 120513 | 3722 | 1669642 |
| -0.045 | 251 | 7 | 0.028 | 244 | 0.972 | 16504482 | 0.078 | 126685 | 3722 | 1103575 |
| -0.0425 | 344 | 9 | 0.026 | 335 | 0.974 | 21029743 | 0.099 | 126604 | 3722 | 1103575 |
| -0.04 | 491 | 34 | 0.069 | 457 | 0.931 | 27425109 | 0.129 | 125085 | 3722 | 1587079 |
| -0.0375 | 600 | 51 | 0.085 | 549 | 0.915 | 32469870 | 0.152 | 127435 | 3722 | 1523317 |
| -0.035 | 714 | 29 | 0.041 | 685 | 0.959 | 40139759 | 0.188 | 129158 | 3722 | 1523317 |
| -0.0325 | 896 | 28 | 0.031 | 868 | 0.969 | 54618153 | 0.256 | 127508 | 3722 | 1523317 |
| -0.03 | 1095 | 27 | 0.025 | 1068 | 0.975 | 58169634 | 0.273 | 124053 | 3722 | 1520567 |
| -0.0275 | 1424 | 50 | 0.035 | 1374 | 0.965 | 71572328 | 0.336 | 129890 | 3722 | 1523317 |
| -0.025 | 1755 | 84 | 0.048 | 1671 | 0.952 | 84622362 | 0.397 | 129186 | 3722 | 1523317 |
| -0.0225 | 2197 | 122 | 0.056 | 2075 | 0.944 | 96539767 | 0.453 | 129974 | 3722 | 1669642 |
| -0.02 | 2712 | 167 | 0.062 | 2545 | 0.938 | 113286288 | 0.532 | 130550 | 3124 | 1669642 |
| -0.0175 | 3432 | 172 | 0.05 | 3260 | 0.95 | 129062034 | 0.606 | 138471 | 3722 | 1669642 |
| -0.015 | 4453 | 195 | 0.044 | 4258 | 0.956 | 144158359 | 0.677 | 139470 | 3177 | 1669642 |
| -0.0125 | 5760 | 250 | 0.043 | 5510 | 0.957 | 162572516 | 0.763 | 143102 | 3124 | 1669642 |
| -0.01 | 7267 | 310 | 0.043 | 6957 | 0.957 | 174258822 | 0.818 | 146984 | 3124 | 1669642 |
| -0.0075 | 9265 | 361 | 0.039 | 8904 | 0.961 | 183906587 | 0.864 | 150587 | 3177 | 1669642 |
| -0.005 | 11486 | 483 | 0.042 | 11003 | 0.958 | 188494233 | 0.885 | 153650 | 3177 | 1669642 |

**Table S6.** chi squared results for number BDMIs per scaffold. X(2) is X(2) analysis threshold. Obs. and Exp. are observed and expected BMDIs. Adjusted p values are Bonferroni corrected for multiple testing.

| **chrom** | **X(2)** | **Obs. BDMI** | **Exp. BDMI** | **chi^2^** | **p (adj.)** |
| --- | --- | --- | --- | --- | --- |
| Scaffold03 | -0.06 | 88 | 146 | 14 | 4.83e-03 |
| Scaffold03 | -0.0575 | 103 | 165 | 14 | 5.02e-03 |
| Scaffold03 | -0.055 | 124 | 187 | 12 | 1.18e-02 |
| Scaffold03 | -0.0525 | 149 | 216 | 12 | 1.48e-02 |
| Scaffold03 | -0.05 | 169 | 248 | 15 | 3.57e-03 |
| Scaffold03 | -0.0475 | 206 | 284 | 12 | 1.35e-02 |
| Scaffold03 | -0.045 | 259 | 326 | 7 | 1.70e-01 |
| Scaffold03 | -0.0425 | 318 | 381 | 6 | 5.10e-01 |
| Scaffold03 | -0.04 | 383 | 445 | 5 | 9.11e-01 |
| Scaffold03 | -0.0375 | 465 | 522 | 3 | 1.00e+00 |
| Scaffold03 | -0.035 | 559 | 614 | 2 | 1.00e+00 |
| Scaffold03 | -0.0325 | 698 | 732 | 1 | 1.00e+00 |
| Scaffold03 | -0.03 | 830 | 885 | 2 | 1.00e+00 |
| Scaffold03 | -0.0275 | 972 | 1073 | 5 | 7.14e-01 |
| Scaffold03 | -0.025 | 1169 | 1310 | 8 | 1.28e-01 |
| Scaffold03 | -0.0225 | 1392 | 1604 | 15 | 2.87e-03 |
| Scaffold03 | -0.02 | 1670 | 1974 | 26 | 1.18e-05 |
| Scaffold03 | -0.0175 | 1995 | 2436 | 44 | 7.26e-10 |
| Scaffold03 | -0.015 | 2454 | 3050 | 66 | 1.49e-14 |
| Scaffold03 | -0.0125 | 3027 | 3894 | 111 | 1.53e-24 |
| Scaffold03 | -0.01 | 3746 | 5122 | 220 | 2.31e-48 |
| Scaffold03 | -0.0075 | 4840 | 6932 | 388 | 7.56e-85 |
| Scaffold03 | -0.005 | 6637 | 9766 | 633 | 2.65e-138 |
| Scaffold17 | -0.06 | 1413 | 264 | 788 | 4.36e-172 |
| Scaffold17 | -0.0575 | 1642 | 297 | 935 | 5.88e-204 |
| Scaffold17 | -0.055 | 1902 | 337 | 1097 | 3.43e-239 |
| Scaffold17 | -0.0525 | 2218 | 389 | 1288 | 1.03e-280 |
| Scaffold17 | -0.05 | 2535 | 446 | 1471 | 2.00e-320 |
| Scaffold17 | -0.0475 | 2871 | 512 | 1655 | 0.00e+00 |
| Scaffold17 | -0.045 | 3212 | 588 | 1824 | 0.00e+00 |
| Scaffold17 | -0.0425 | 3637 | 687 | 2028 | 0.00e+00 |
| Scaffold17 | -0.04 | 4102 | 802 | 2241 | 0.00e+00 |
| Scaffold17 | -0.0375 | 4594 | 941 | 2436 | 0.00e+00 |
| Scaffold17 | -0.035 | 5126 | 1106 | 2624 | 0.00e+00 |
| Scaffold17 | -0.0325 | 5712 | 1320 | 2780 | 0.00e+00 |
| Scaffold17 | -0.03 | 6519 | 1595 | 3035 | 0.00e+00 |
| Scaffold17 | -0.0275 | 7403 | 1934 | 3262 | 0.00e+00 |
| Scaffold17 | -0.025 | 8473 | 2362 | 3520 | 0.00e+00 |
| Scaffold17 | -0.0225 | 9866 | 2891 | 3910 | 0.00e+00 |
| Scaffold17 | -0.02 | 11646 | 3558 | 4434 | 0.00e+00 |
| Scaffold17 | -0.0175 | 13790 | 4393 | 5035 | 0.00e+00 |
| Scaffold17 | -0.015 | 16529 | 5499 | 5771 | 0.00e+00 |
| Scaffold17 | -0.0125 | 20247 | 7021 | 6777 | 0.00e+00 |
| Scaffold17 | -0.01 | 25494 | 9234 | 8169 | 0.00e+00 |
| Scaffold17 | -0.0075 | 31628 | 12497 | 9080 | 0.00e+00 |
| Scaffold17 | -0.005 | 39133 | 17606 | 9190 | 0.00e+00 |

**Table S7.** Wilcox rank sum test results for distance to centromeres. P values are corrected for multiple testing (Bonferroni correction) within each analysis.

| **Analysis** | **Obs. median** | **Boot. median** | **W** | **p** | **p (adj.)** |
| --- | --- | --- | --- | --- | --- |
| gIMble | 2418184 | 42287 | 12383464 | 2.48e-12 | 2.48e-12 |
| X(2) -0.06 | 2809237 | 4220426 | 1104844 | 1.12e-07 | 2.57e-06 |
| X(2) -0.0575 | 2311584 | 4360588 | 942392 | 3.37e-09 | 7.75e-08 |
| X(2) -0.055 | 2316322 | 4192145.5 | 856228 | 7.58e-08 | 1.74e-06 |
| X(2) -0.0525 | 2878366 | 4459107.5 | 1556164 | 6.31e-08 | 1.45e-06 |
| X(2) -0.05 | 2609456 | 4617692.5 | 4104332 | 5.86e-15 | 1.35e-13 |
| X(2) -0.0475 | 2311584 | 4466590.5 | 9066680 | 2.59e-23 | 5.96e-22 |
| X(2) -0.045 | 1044832 | 4163179 | 17316882 | 2.10e-47 | 4.83e-46 |
| X(2) -0.0425 | 688747 | 4018955 | 35227564 | 7.02e-79 | 1.61e-77 |
| X(2) -0.04 | 793272 | 3918297.5 | 71252232 | 9.07e-88 | 2.09e-86 |
| X(2) -0.0375 | 1071584 | 4031945 | 88890126 | 1.69e-102 | 3.88e-101 |
| X(2) -0.035 | 1833192 | 4027233.5 | 152834358 | 5.46e-98 | 1.26e-96 |
| X(2) -0.0325 | 2294802 | 4215404 | 259332734 | 5.53e-98 | 1.27e-96 |
| X(2) -0.03 | 2621358 | 4154846.5 | 356180290 | 4.33e-92 | 9.97e-91 |
| X(2) -0.0275 | 2579866 | 4209004.5 | 652446318 | 1.07e-132 | 2.45e-131 |
| X(2) -0.025 | 2579866 | 4239258 | 1012827258 | 1.02e-165 | 2.35e-164 |
| X(2) -0.0225 | 2579866 | 4210440.5 | 1931505090 | 5.18e-212 | 1.19e-210 |
| X(2) -0.02 | 2345526 | 4241318.5 | 2914818130 | 2.04e-293 | 4.68e-292 |
| X(2) -0.0175 | 2379454 | 4249533 | 4674881678 | 0.00e+00 | 0.00e+00 |
| X(2) -0.015 | 2463600 | 4323157.5 | 7932668552 | 0.00e+00 | 0.00e+00 |
| X(2) -0.0125 | 2291478 | 4335050 | 1,3015E+10 | 0.00e+00 | 0.00e+00 |
| X(2) -0.01 | 2394696 | 4350845.5 | 2,2989E+10 | 0.00e+00 | 0.00e+00 |
| X(2) -0.0075 | 2290832 | 4400961 | 3,9541E+10 | 0.00e+00 | 0.00e+00 |
| X(2) -0.005 | 2294905 | 4409323 | 6,3385E+10 | 0.00e+00 | 0.00e+00 |

**Table S8.** Coverage and overlap between long-term gIMble barriers and candidate BDMIs.

| **x2_cutoff** | **AD_total_cov** | **AD_gim_over** | **gim_total_cov** | **bp_dmi_only** | **bp_gim_only** | **perc_dmi_only** | **perc_gim_only** | **perc_both** |
| --- | --- | --- | --- | --- | --- | --- | --- | --- |
| -0.06 | 4250 | 535 | 17442 | 3715 | 16908 | 0.18 | 0.8 | 0.03 |
| -0.0575 | 4975 | 598 | 17442 | 4377 | 16845 | 0.2 | 0.77 | 0.03 |
| -0.055 | 5046 | 561 | 17442 | 4485 | 16881 | 0.2 | 0.77 | 0.03 |
| -0.0525 | 7989 | 671 | 17442 | 7318 | 16772 | 0.3 | 0.68 | 0.03 |
| -0.05 | 8190 | 755 | 17442 | 7436 | 16688 | 0.3 | 0.67 | 0.03 |
| -0.0475 | 12906 | 963 | 17442 | 11943 | 16479 | 0.41 | 0.56 | 0.03 |
| -0.045 | 16504 | 1240 | 17442 | 15265 | 16203 | 0.47 | 0.5 | 0.04 |
| -0.0425 | 21030 | 1678 | 17442 | 19352 | 15764 | 0.53 | 0.43 | 0.05 |
| -0.04 | 27425 | 1907 | 17442 | 25518 | 15535 | 0.59 | 0.36 | 0.04 |
| -0.0375 | 32470 | 2793 | 17442 | 29677 | 14649 | 0.63 | 0.31 | 0.06 |
| -0.035 | 40140 | 4073 | 17442 | 36067 | 13369 | 0.67 | 0.25 | 0.08 |
| -0.0325 | 54618 | 4952 | 17442 | 49666 | 12491 | 0.74 | 0.19 | 0.07 |
| -0.03 | 58170 | 5467 | 17442 | 52702 | 11975 | 0.75 | 0.17 | 0.08 |
| -0.0275 | 71572 | 7016 | 17442 | 64556 | 10426 | 0.79 | 0.13 | 0.09 |
| -0.025 | 84622 | 7708 | 17442 | 76914 | 9735 | 0.82 | 0.1 | 0.08 |
| -0.0225 | 96540 | 8757 | 17442 | 87783 | 8686 | 0.83 | 0.08 | 0.08 |
| -0.02 | 113286 | 10121 | 17442 | 103165 | 7321 | 0.86 | 0.06 | 0.08 |
| -0.0175 | 129062 | 11901 | 17442 | 117162 | 5542 | 0.87 | 0.04 | 0.09 |
| -0.015 | 144158 | 12792 | 17442 | 131366 | 4650 | 0.88 | 0.03 | 0.09 |
| -0.0125 | 162573 | 14441 | 17442 | 148132 | 3002 | 0.89 | 0.02 | 0.09 |
| -0.01 | 174259 | 15371 | 17442 | 158887 | 2071 | 0.9 | 0.01 | 0.09 |
| -0.0075 | 183907 | 15869 | 17442 | 168037 | 1573 | 0.91 | 0.01 | 0.09 |
| -0.005 | 188494 | 16235 | 17442 | 172260 | 1208 | 0.91 | 0.01 | 0.09 |

**Table S9**. List of 13 genes identified in regions with persistent BDMIs exhibiting no gene flow between *F. aquilonia* and *F. polyctena*.

| **Gene ID** | **Best BLAST hit** | **E-value** | **Identity (%)** | **Prediction** |
| --- | --- | --- | --- | --- |
| jg6572.t1 | XM_029819064.1 | 3.88E-162 | 81.51 | Formica exsecta uncharacterized LOC115242631 (LOC115242631) |
| jg6572.t1 | XM_029802978.1 | 2.20E-146 | 95.32 | Formica exsecta uncharacterized LOC115232845 (LOC115232845) |
| jg6573.t1 | XM_029802977.1 | 3.70E-149 | 99.09 | Formica exsecta protein G12-like (LOC115232844) |
| jg6574.t1 | XM_029802909.1 | 1.53E-156 | 98.25 | Formica exsecta uncharacterized LOC115232803 (LOC115232803) |
| jg6574.t1 | XM_050591825.1 | 1.09E-131 | 85.02 | Cataglyphis hispanica protein G12-like (LOC126849703) |
| jg20385.t1 | XM_029804083.1 | 8.17E-22 | 100.00 | Formica exsecta proteoglycan 4-like (LOC115233581), transcript variant X2 |
| jg20385.t1 | XM_029804084.1 | 8.17E-22 | 100.00 | Formica exsecta proteoglycan 4-like (LOC115233581), transcript variant X3 |
| jg20385.t1 | XM_029804081.1 | 8.17E-22 | 100.00 | Formica exsecta proteoglycan 4-like (LOC115233581), transcript variant X1 |
| jg20386.t1 | XM_029804080.1 | 0.00E+00 | 99.26 | Formica exsecta ras-related protein Rab-34 (LOC115233580) |
| jg20387.t1 | XM_029804088.1 | 5.82E-116 | 100.00 | Formica exsecta 60S ribosomal protein L23a (LOC115233585) |
| jg20388.t1 | XM_029804043.1 | 0 | 99.17 | Formica exsecta WASH complex subunit 4 (LOC115233553), transcript variant X3 |
| jg20388.t1 | XM_029804042.1 | 0 | 99.17 | Formica exsecta WASH complex subunit 4 (LOC115233553), transcript variant X2 |
| jg20388.t1 | XM_029804041.1 | 0 | 99.17 | Formica exsecta WASH complex subunit 4 (LOC115233553), transcript variant X1 |
| jg20389.t1 | XM_029804044.1 | 1.34E-163 | 99.57 | Formica exsecta CUE domain-containing protein 1 (LOC115233554), transcript variant X1 |
| jg20390.t1 | XM_050601544.1 | 0 | 97.95 | Cataglyphis hispanica ethanolaminephosphotransferase 1-like (LOC126854617) |
| jg20391.t1 | XM_029804033.1 | 1.94E-65 | 99.00 | Formica exsecta coiled-coil-helix-coiled-coil-helix domain-containing protein 7 (LOC115233546) |
| jg20392.t1 | XM_029804032.1 | 5.49E-162 | 99.59 | Formica exsecta exosome complex component RRP40 (LOC115233545) |
| jg20945.t1 | XM_029821704.1 | 2.21E-59 | 92.31 | Formica exsecta uncharacterized LOC115244228 (LOC115244228), transcript variant X1 |
| jg12768.t1 | XM_025408744.1 | 5.52E-165 | 85.05 | Camponotus floridanus uncharacterized LOC112637969 (LOC112637969) |
